# Symphonizing pileup and full-alignment for deep learning-based long-read variant calling

**DOI:** 10.1101/2021.12.29.474431

**Authors:** Zhenxian Zheng, Shumin Li, Junhao Su, Amy Wing-Sze Leung, Tak-Wah Lam, Ruibang Luo

## Abstract

Deep learning-based variant callers are becoming the standard and have achieved superior SNP calling performance using long reads. In this paper, we present Clair3, which leveraged the best of two major method categories: pile-up calling handles most variant candidates with speed, and full-alignment tackles complicated candidates to maximize precision and recall. Clair3 ran faster than any of the other state-of-the-art variant callers and performed the best, especially at lower coverage.

## Maintext

The first preprint of DeepVariant^1^ was released in late 2016, marking the beginning of the use of deep learning-based methods (DL methods) instead of traditional statistical methods for variant calling. Over the years, several DL methods have been developed. We are now witnessing a complete take-over, led by DeepVariant for short-read variant calling. Long-read variant calling, using Oxford Nanopore (ONT) data, on the other hand, has been dominated by DL-methods since the beginning, primarily owing to the difficulty caused by its higher base error rate in general. Although the DL methods for short-read and long-read have a lot in common, the problem of long-read variant calling is considered more difficult. This led to two major designs – using pileup or full-alignment as the input of the decision-making neural network – which are significantly different in both performance and speed.

Long-read variant callers, including Clairvoyante^2^, Clair^3^, and Nanocaller^4^, are pileup-based, in which the read alignments are summarized into features and counts before being inputted into a variant calling network. PEPPER-Margin-DeepVariant^5^ (PEPPER) is full alignment-based. The input to the DeepVariant variant calling network is kept with spatial information in the read alignments and is tens of times larger than the pileup inputs in terms of size. Medaka^6^ is consensus-based; it uses pileup input to generate a diploid consensus in the first iteration and two haploid consensuses in the second. The differences between the reference and consensuses are identified and combined into variants. NanoCaller uses a 3-layer CNN architecture, which is similar to the architecture of our first-generation caller Clairvoyante. These are all state-of-the-art algorithms; the pileup-based algorithms are usually superior in terms of time efficiency and the full-alignment algorithms provide the best precision and recall. More characteristics and limitations of the two designs are discussed in Clair^3^ and DeepVariant^1^, respectively. However, while the two designs are not mutually exclusive, there have not been any studies combining pileup calling and full-alignment calling.

To fill the gap, we developed Clair3, the successor to Clair, which makes the best of both designs. It runs as fast as the pileup-based callers and performs as well as the full alignment-based callers. **Extended Data Figure 1** shows the workflow for Clair3. The philosophy behind Clair3 is to trust the full-alignment model unless the pileup model can make a quick but reliable decision. First, the pileup calling network goes through all the variant candidates that passed a coverage threshold and an alternative allele frequency threshold. Next, the high-quality pileup calls are used to phase the alignments and as part of the final output. Then, the alignments phased by WhatsHap^7^ are used to generate full-alignment input that is ∼23 times larger in size than the pileup input for each low-quality pileup call for full-alignment calling. Finally, the full-alignment calls are integrated with the high-quality pileup calls as the final output. More details and parameters about the Clair3 workflow, input/output, and network architecture are provided in **Methods**.

We benchmarked Clair3 v0.1-r11 against PEPPER r0.8, Medaka v1.4.4 (that last version of ONT’s in-house tool that supported variant calling), Longshot^8^ v0.4.5 (non-deep learning-based; works only with SNP), and Clair v2.1.1 (the Clair3 predecessor) on two GIAB^9, 10^ samples: HG003 and HG004. HG003 was tested on models (including a pileup and a full-alignment model) trained on HG001, 2, 4 and 5. HG004 was tested on models trained on HG001, 2, 3 and 5. The model availability and training details are in **Methods**. We primarily benchmarked ONT data base-called using Guppy 5 (version 5.0.14) data, but in addition we also benchmarked Guppy 4 (version 4.2.2) for two reasons: 1) variant caller’s capability can be shown on noisier data - compared to the Guppy 5, which was released in mid-2021, Guppy 4’s read accuracy is ∼1.8% lower^11^, so it can better demonstrate the speed and performance of different variant calling methods, and 2) data compatibility - as at the completion date of this paper, Guppy 4 base-called reads were still the latest version available for download by the Human Pangenome Reference Consortium^12^. A summary of the datasets used for training and testing is shown in **Supplementary Table 1**. The correct PEPPER and Medaka models for either Guppy 5 or 4 data were used for benchmarking. The links to the dataset, and the versions, commands and parameters used for each tool are available in the **Supplementary Notes**.

The benchmarking results at coverage from 10x to 50x of Guppy 5 data are shown in **Figure 1a, Supplementary Table 2**, and **Supplementary Table 3**. The observations of different tools on HG003 and HG004 are almost identical, ruling out the possibility of any tools’ overfitting to a particular sample. In terms of the SNP F1-score, Clair3 outperformed all other tools at the more challenging lower coverage (10x to 20x). Above 20x, Clair3 performed similar to PEPPER above 20x, but ran much faster. Different from the SNP F1-score that plateaued above 20x, the Indel F1-score kept increasing with coverage. Looking at the precision and recall at 20x, which is an optimistic estimation of the minimal coverage achievable per PromethION flowcell in production, in terms of SNP (**Figure 1b**), Clair3 achieved 99.22% and 99.42%, compared to PEPPER’s 99.66% and 98.99%, in HG003. In terms of Indel, Clair3 achieved 88.56% and 62.33%, compared to PEPPER’s 86.06% and 63.26%. In terms of speed (**Figure 1c**), at 20x, Clair3 and Clair ran the fastest (∼5 hours). PEPPER ran about four times slower than Clair3 (∼20 hours). We then benchmarked using the noisier and more challenging Guppy 4 data, on which Clair3 has more significantly outperformed the other tools (**Supplementary Table 4** and **Supplementary Table 5**). Using Guppy 4 data, Clair3 significantly outperformed the other tools from 10x to 30x. At 20x in HG003, in terms of SNP, Clair3 achieved 99.17% and 98.77%, compared to PEPPER’s 90.40% and 98.95%. In terms of Indel, Clair3 achieved 80.91% and 53.33%, compared to PEPPER’s 73.73% and 48.53%. Using Guppy 4 data, Clair3 ran about three times faster than PEPPER (∼8 vs. ∼30 hours). To observe the performance characteristics of Clair3 at different genomic contexts, we compared Clair3 to PEPPER according to the GIAB genome stratifications^13^ v2.0 on HG003 at sufficient coverage of (50x) of the noisier Guppy 4 data. The results are shown in **Supplementary Figure 1** and **Supplementary Table 6**. In SNPs, Clair3 outperformed PEPPER on precision in low complexity and functional regions. Clair3 and PEPPER had the same recall in different regions. In Indels, Clair3 outperformed PEPPER in both precision and recall in all regions.

**Figure 1.**
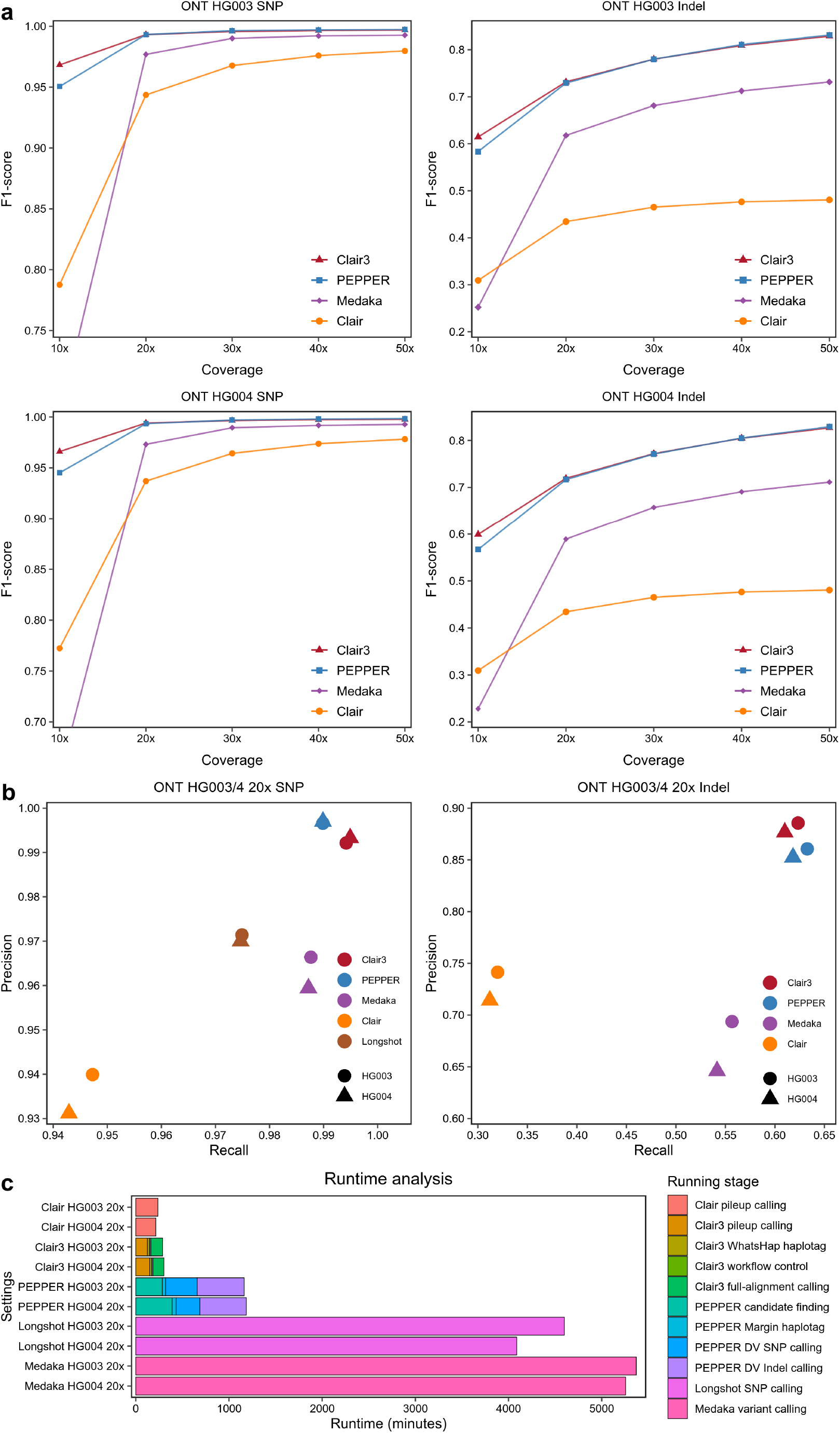
Benchmarking results on HG003 and HG004 with Guppy 5 data. (a) The SNP/Indel F1-score of different tools at multiple coverage from 10x to 50x. In terms of the SNP F1-score, Clair3 outperformed the other tools at the more challenging lower coverage (10x to 20x). (b) The precision against the recall of different tools at 20x coverage. (c) The runtime breakdowns of different tools at 20x coverage. Clair3 and Clair ran the fastest (∼5 hours), PEPPER ran about four times slower than Clair3 (∼20 hours).

The success of the Clair3 method lies in the effective distinction between true and false calls during pileup calling, so that only necessary candidates are sent to the much more computationally intensive full-alignment calling. **Figure 2a** shows that an effective distinction was achieved using variant quality. Using HG003 at 50x as an example, most false variant calls and false reference calls had a quality between 0 to 10, while the true calls were between 15 to 30. In reality, while the correctness of a pileup call is not known in advance, we empirically decided to send the bottom 30% of the pileup variant calls and the bottom 10% of the pileup reference calls to full-alignment calling as the default settings of Clair3 (See **Methods**). In the previous example, quality cut-off 16 was chosen for the variant calls, which resulted in 96% of the false variant calls and only 7% of the true variant calls being sent to full-alignment calling. Similarly, cut-off 16 was chosen for the reference calls, so that 98% of the false reference calls and only 13% of the true reference calls were sent to full-alignment calling. **Figure 2b** shows that ∼60% of the pileup failed variant calls and ∼30% of the pileup failed reference calls were correctly called in full-alignment calling. We tested sending different percentages of pileup variant calls to full-alignment calling, from 0% (pileup calling only) to 100% (full-alignment calling only). The results are shown in **Figure 2c** and **Supplementary Table 8**. Clair3’s default, which had a similar performance to full-alignment calling but ran ∼3 times faster, showed that integrating pileup and full-alignment calling is a better strategy than relying on only one of them. To eliminate the concern of model overfitting, we benchmarked chromosome 20, which has always been excluded from model training. The results shown in Supplementary Table 7 concluded that no overfitting is observed in Clair3.

**Figure 2.**
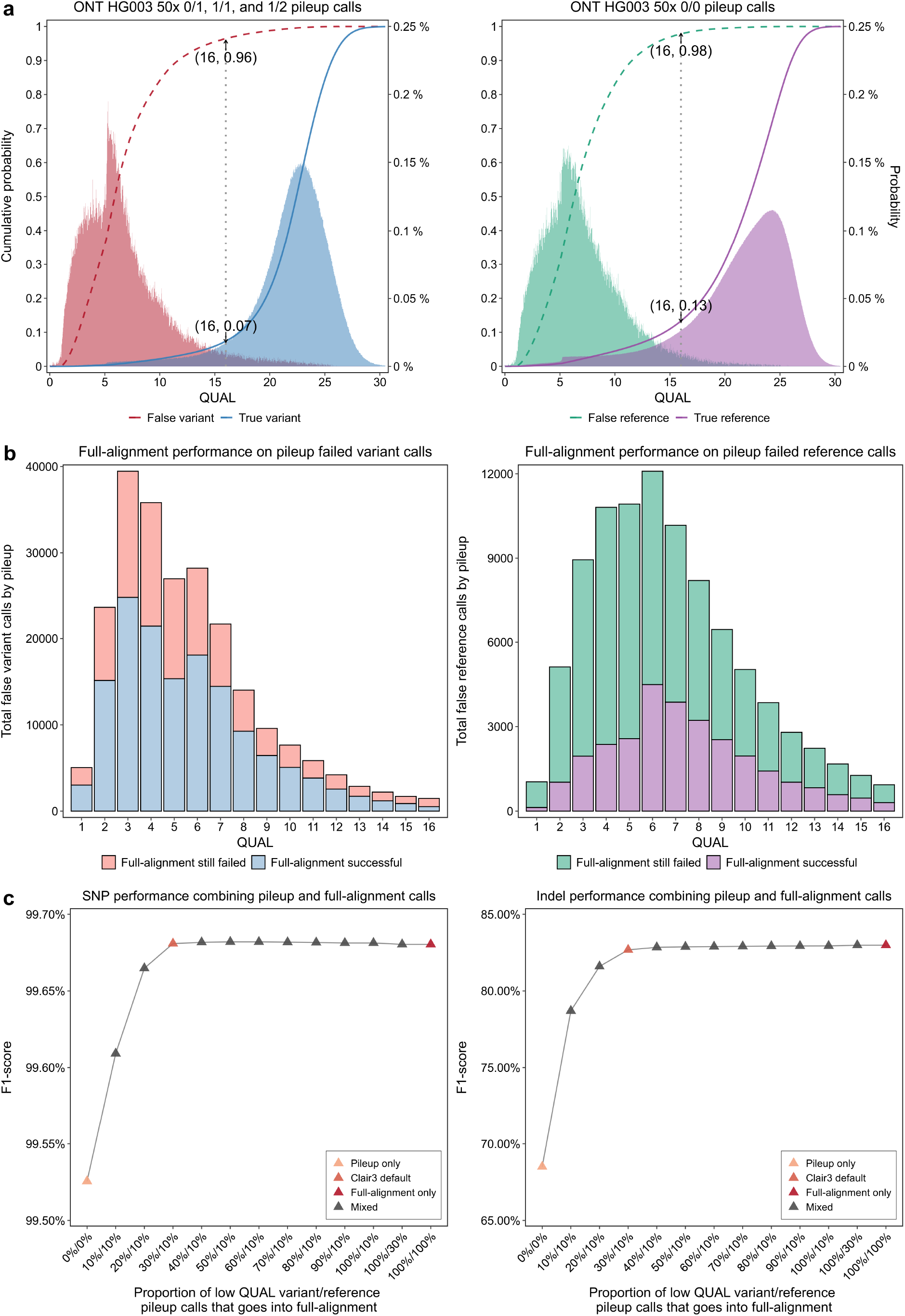
Pileup and full-alignment calling working details and synergy on HG003 at 50x coverage of Guppy 5 data. (a) The variant quality distribution of the true and false variant/reference pileup calls. The figure shows that an effective distinction was achieved using variant quality. (b) The performance of full-alignment on pileup failed variants of different variant quality. The figure shows that ∼60% of the pileup failed variant calls and ∼30% of the pileup failed reference calls were correctly called in full-alignment calling. (c) The F1-score when different proportions of low-quality variant/reference calls enter full-alignment calling. QUAL is the variant quality as defined in the VCF specifications. The figure shows that integrating pileup and full-alignment calling is a better strategy than relying on only one of them.

The benchmarks focused on the more challenging ONT data, but the Clair3 method is not restricted to a particular sequencing technology. It should work particularly well in terms of both runtime and performance on noisy data. Having integrated plenty of feedback from the community and ONT, Clair3 is currently in its eleventh revision. We observed that ONT has removed the variant calling submodule from Medaka and suggested Clair3 for variant calling since v1.5.0^14^. We also observed that in PEPPER’s r0.7^15^ update, a module in the front of the pipeline that was used solely for variant candidate selection was repurposed to output summary-based variant calls to relieve the heavy full-alignment calling workload, which is converging to Clair3’s idea. We expect integrating pileup and full-alignment calling to be a common practice in deep learning-based variant calling in the future.

## Method

### The Clair3 workflow

As **Extended Data Figure 1** shows, pileup candidates that are above a coverage threshold and an allele frequency threshold are extracted, and then called using the pileup network. The pileup calls are grouped into variant calls (genotype 0/1, 1/1, and 1/2) and reference calls (0/0). Both groups are ranked according to variant quality (QUAL). High-quality heterozygous SNP calls (top 70% in 0/1 calls) are used in WhatsHap phasing to produce phased alignment for input to the full-alignment network. Low-quality pileup calls (defaulted to the lowest 30% of variants and 10% of reference calls) are then called again using the full-alignment network. Finally, the full-alignment calls and high-quality pileup calls are outputted. Clair3 supports both VCF and GVCF output formats.

### Input/Output

Clair3 uses a pileup input design simplified from that of its predecessors, and a full-alignment input to cover as many details in the read alignments as possible. **Supplementary Figure 2** visualizes the pileup and full-alignment inputs of a random SNP, insertion, deletion, or non-variant. **The pileup input** is 594 integers – 33 genome positions wide with 18 features at each position – A+, C+, G+, T+, I_S_+, I^1^_S_+, D_S_+, D^1^_S_+, D_R_+, A-, C-, G-, T-, I_S_-, I^1^_S_-, D_S_-, D^1^_S_-, and D_R_-. +, - means the positive strand and negative strand. A, C, G, T, I, D means the count of read support of the four nucleotides, insertion, and deletion. Superscript “1” means only the indel with the highest read support is counted if various lengths of indel were found in a candidate site. (i.e., all indels are counted if without “1”). Subscript “S” means the starting position of an indel. Subscript “R” means the following positions of an indel. For example, a 3bp deletion with the most reads support will have the first deleted base counted in either D^1^_S_+ or D^1^_S_-, and the second and third deleted bases counted in either D_R_+ or D_R_-. The design was determined experimentally, but the rationale is that for 1bp indels that are easy to call, look into the differences between the “S” counts, but reduce the quality if the “R” counts and discrepancy between positions increase. Supplementary Figure 2 provides some intuitions on how the features are counted given four random examples. For developers to confirm their understanding, the input creation logics are available at https://github.com/HKU-BAL/Clair3/blob/main/preprocess/CreateTensorPileup.py. **The pileup output** is explained in the “Network outputs of the pileup and full-alignment network” section in the Supplementary Materials. The indel allele (or two indel alleles) with the highest reads support is used as the output according to the decision made in the 21-genotype task. **The full-alignment input** is 23,496 integers – 8 channels of 33 genome positions and 89 maximum number of reads. The description of the eight channels is in the Supplementary Note. **The full-alignment output** is explained in the “Network outputs of the pileup and full-alignment network” section in the Supplementary Materials. The two indel length tasks can represent the exact indel length from -15 to 15bp, or below -15bp/above 15bp. An indel call with an exact length will output the most reads-supported allele at that length. Otherwise, the most reads-supported allele below -15bp/above 15bp is outputted. In training, indel length task 1 is given the smaller number, and in all our variant calling experiments, no length predictions in task 1 larger than in task 2 were observed. **The maximum supported coverage** of full-alignment input was 89. If the coverage was above 89, random subsampling of reads was applied. If the coverage was below 89, zero-padding was applied with reads placed at the center of the input. The maximum supported coverage of full-alignment input can be increased by changing the “matrix_depth_dict” variable in the “param_f.py” configuration file.

### Network architecture

The pileup and full-alignment networks are shown in **Supplementary Figure 3. The pileup network** uses two bidirectional long short-term memory (Bi-LSTM) layers with 128 and 160 LSTM units. Stacked LSTM layers enable the network to learn the characteristics of raw sequential signal from different aspects at each position, but without increasing memory capacity, which enables the network to converge faster. Compared to Clair, the transpose-split layer is removed for a 40% speedup with a small performance loss that is taken care of in full-alignment calling. **The full-alignment network** is derived from residual neural network (ResNet) and uses three standard residual blocks. A convolutional layer is added on top of each residual block to expand channels but reduce dimensionality across channels. A spatial pyramid pooling^16^ (SPP) layer is used to tackle the problem of varying coverage in full-alignment input. SPP is a pooling layer that removes a network’s fixed-size constraint, thus avoiding the need for input cropping or warping at the beginning. The SPP layer generates various receptive fields using three pooling scales (1×1, 2×2, and 3×3) in each channel. It then pools the receptive fields of all channels and generates a fixed-length output for the next layer. In both networks, the dropout rates of 0.2 for the flatten layer, 0.5 for the penultimate dense layer, and 0.2 for the task-specific final dense layers, are empirically determined. In comparison, the Inception-v3 network used as full-alignment network in DeepVariant and PEPPER is ∼8 times larger (2,989,210 vs. ∼24 million parameters) than Clair3’s full-alignment network.

We tried removing a residual block from the full-alignment network, the overall F1-score reduced by ∼4% in average in multiple experiments with HG003 and HG004, and coverage from 10x to 50x. Adding a residual block, the overall F1-score improvements were unnoticeable, but the network speed slowed down by 20%. Removing the SPP layer reduced the Indel F1-score by ∼10% at 10x coverage. More results of removing the insertion, phasing, MQ, BQ channel, or the two Indel length tasks are shown in the **Supplementary Table 9**. A visualization toolkit that shows the network activations of individual inputs using guided propagation is available at the GitHub repository.

### Model availability and training

Pretrained models are provided in Clair3’s installation. Models for specific chemistries and basecallers that are tested and supported by the ONT developers are available through Rerio (https://github.com/nanoporetech/rerio). The detailed steps, options, and caveats for training a pileup model and a full-alignment model are available in Clair3’s GitHub repo (at https://github.com/HKU-BAL/Clair3/blob/main/docs/pileup_training.md and https://github.com/HKU-BAL/Clair3/blob/main/docs/full_alignment_training_r1.md) and are continually updated. The pretrained models, while targeted for use in production, were trained using multiple GIAB samples with known variants and 10 coverages for each sample (more details in the “Training data augmentation using subsampled coverage” section in Supplementary Materials), but they always hold out chromosome 20 in Clair3. We used the following new training technics in Clair3. **(1) Representation Unification**: a variant can be represented in multiple forms^13^. Traditional variant calling methods rely on postprocessing (e.g., hap.py, RTG Tools) to equate multiple forms. However, to generate correct training samples, Clair3 must unify a variant’s representations between the alignments and the truth variants. **Supplementary Figure 4** shows four cases in which the alignments and the truth variants have different representations that would confuse the training if not unified. Clair3 chooses to align the truth variants’ representation to the alignments. The five detailed steps are shown in **Supplementary Figure 5**. First, the truth variants and alignments are phased (if not yet done) using WhatsHap. Second, among the candidates with alternative allele frequency ≥0.15, confident and *in situ* matches between the alignments and truth variants are identified and excluded from computationally intensive step 3. Third, the best match between the possible haplotypes of the truth variants and candidates is sought. Each of the truth variants can be either positive (using its reported genotype) or negative (using 0|0), and their Cartesian product forms possible haplotypes of the truth variants. Similarly, each candidate can be either 0|0, 0|1 (or 1|0 according to the phased alignments), or 1|1, and their Cartesian product forms the possible haplotypes of the candidates. A pairwise comparison is then done to find equivalent haplotypes between the two Cartesian products, and among all equivalents, the candidate haplotype with the most reads support is selected. The variants in the haplotype are used as the new truth variants. This step is computationally intensive, so in practice, we applied the step to partitions with at most 15 candidates and required less than 100bp between the candidates. Fourth, low alternative allele frequency (≥0.08 but <0.15) candidates with *in situ* matches between the alignments and the truth variants were chosen. Fifth, the truth variants or unified variants generated in steps 2, 3 and 4 were merged. In our benchmarks, representation unification alone in general increased the SNP recall by ∼0.2% and Indel recall by ∼2%. **(2) Ratio of variants to non-variants samples for training**: In Clair, the ratio was fixed at 1:2. In Clair3, we tested ratios up to 1:10 for both pileup and full-alignment model training, and we observed a monotonic but decelerated performance increase with more non-variants added to the training. Since focal loss is used to alleviate the effect of training class imbalance, another possible explanation is that the 21-genotype output task that Clair3 relies primarily on is insensitive to the ratio because it judges only the genotype of a candidate instead of whether a candidate is a variant or not. We chose 1:5 and 1:1 as the default ratio for pileup and full-alignment model training, respectively, to strike a balance between model performance and training speed. **(3) Use of phased alignments**: Deep-learning and full-alignment based variant callers DeepVariant and PEPPER concluded that using phased alignments is essential to their high performance. In Clair3, high-quality heterozygous pileup calls are used to phase the input alignments using the ‘phase’ and ‘haplotag’ modules in WhatsHap. The phased alignments are used as input for full-alignment calling. When training a full-alignment model, two training samples for each variant, one using phased alignments and the other unphased, are used to ensure the model works when alignments cannot be properly phased. In our benchmarks, the use of phased alignments alone, in general, increased the SNP F1-score by ∼0.1%, and the Indel F1-score by ∼6%. **(4) New optimization methods**: Clair3 removed both the cyclical learning rate and learning rate decay strategies used in Clair, and now uses the Ranger optimizer (RectifiedAdam^17^ plus Lookahead^18^) for automated warm-up, faster convergence, minimal computational overhead, etc. In our benchmarks, compared to Clair, the new optimizer alone, in general, increased the overall F1-score of pileup calling by ∼0.2% (tested with three repetitions with random seed changed in weight initialization).

### Benchmarking methods and computational concerns

We used five GIAB samples, HG001 to 5, for either model training or testing. Following the PrecisionFDA v2 practices^9, 10^, we used HG003 and HG004 for testing. When using either HG003 or HG004 for testing, the other four samples were used for training. We selected 10% of the training samples for validation and chose the best-performing epoch in the first 30 epochs in the validation data for benchmarking. We used hap.py^13^ to compare the called variants against the true variants, and used Clair3’s ‘GetOverallMetrics’ module to generate three metrics, ‘precision’, ‘recall’, and ‘F1-score’, for five categories: ‘overall’, ‘SNP’, ‘Indel’, ‘Insertion’, and ‘Deletion’. The evaluation metrics also followed the PrecisionFDA v2 practices and are further explained in Supplementary Materials. We used qfy.py with V2.0 GIAB genome stratifications to evaluate Clair3’s performance in challenging and targeted regions of the genome. Runtimes and memory consumptions were gauged on a server with two 2.1GHz Intel Xeon Silver 4116s, with 24 cores, and 256GB memory at 2666MHz. With the same setting, Clair3 finished in ∼6 hours using ∼20x of ONT Guppy 4 data and in ∼2 hours with the same amount of Guppy 5 data. The peak/average memory consumption of Clair3 and other tools are shown in **Supplementary Table 10**.

### Brief summary of methods tested showing no or negligible improvement

**(1) Use of more residual blocks in the full-alignment network**: We added a fourth residual block with 512 channels. The number of parameters increased from 2,989,210 to 9,812,634. The runtime doubled, but the performance change was negligible, even though the terminal training loss fell. **(2) Local realignment**: This technique is essential for high indel calling performance in state-of-the-art, short-read, small variant callers. But it worked differently on long reads. We tried local realignment using a 2000bp window in regions with a high density of candidates using a local realignment algorithm similar to that of DeepVariant. We observed that while it increased the recall, local realignment tripled the runtime and introduced ∼10% of new non-variant candidates, which in turn, lowered the precision. In Clair3, we implemented local realignment, but disabled it on long reads as the default. **(3) Including variants outside high-confidence regions in training**: To increase variant training samples, we explored including variants outside the high-confidence regions in training, but observed negative performance improvement in Clair. In Clair3, the GIAB truth datasets we used were upgraded from version 3.3.2 to 4.2.1, but we had the same observation that including variants outside the high-confidence regions in training jeopardized model performance. **(4) Selecting candidates for full-alignment calling based on reference sequence complexity**: Variant calling is more difficult in the “low complexity” and “difficult to map” regions. In addition to selecting candidates by pileup calling quality ranking for full-alignment calling, we added those candidates at positions with relatively low sequence entropy (the lowest 30% of the whole genome). About three times more candidates were selected for full-alignment calling, but the performance increase was negligible.

## Code availability

Clair3 is open-source software (BSD 3-Clause license), hosted by GitHub at https://github.com/HKU-BAL/Clair3, and available through Docker, Bioconda, and Singularity. Clair3 is also available in Zenodo at DOI “10.5281/zenodo.6637001”.

## Data availability

The 1) links to the reference genomes, truth variants, benchmarking materials, and ONT data, and 2) the commands and parameters used in this study, are available in the Supplementary Notes. All analysis output, including the VCFs and running logs, are available at http://www.bio8.cs.hku.hk/clair3/analysis_result.

## Acknowledgements

R. L. was supported by Hong Kong Research Grants Council grants GRF (17113721), ECS (27204518), and TRS (T21-705/20-N and T12-703/19-R), and the URC fund at HKU.

## Author contributions

R. L. conceived the study. Z. Z. and R. L. designed the algorithms, implemented Clair3, and wrote the paper. All authors evaluated the results and revised the manuscript.

---

**Extended Data Figure 1.**
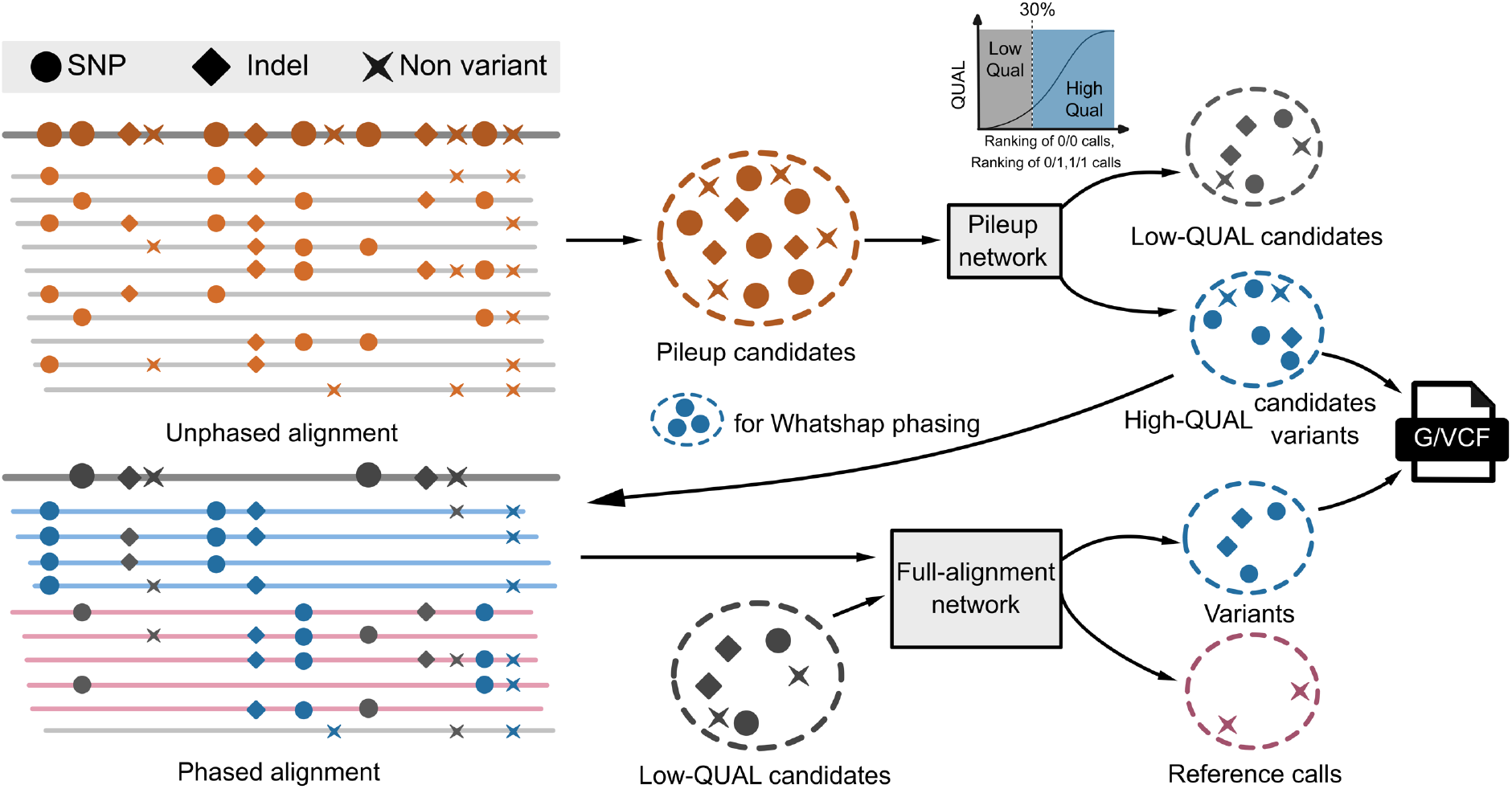
The workflow for Clair3.

