## Supplementary Materials for "Symphonizing pileup and full-alignment for deep learning-based long-read variant calling"

### Supplementary Figures

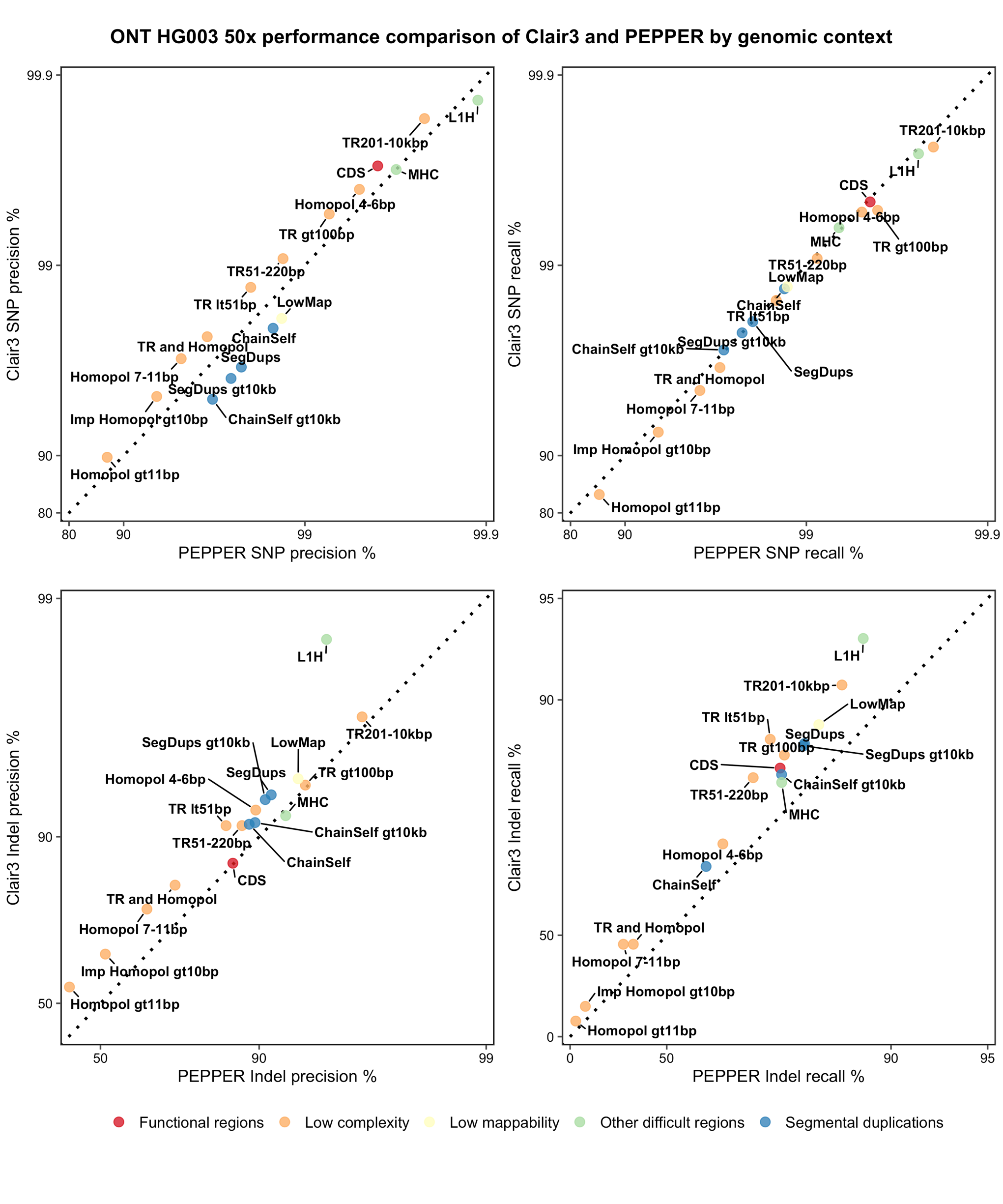

#### **Supplementary Figure 1.** Performance comparison of Clair3 and PEPPER by genomic context on HG003 at 50x of Guppy 4 data.

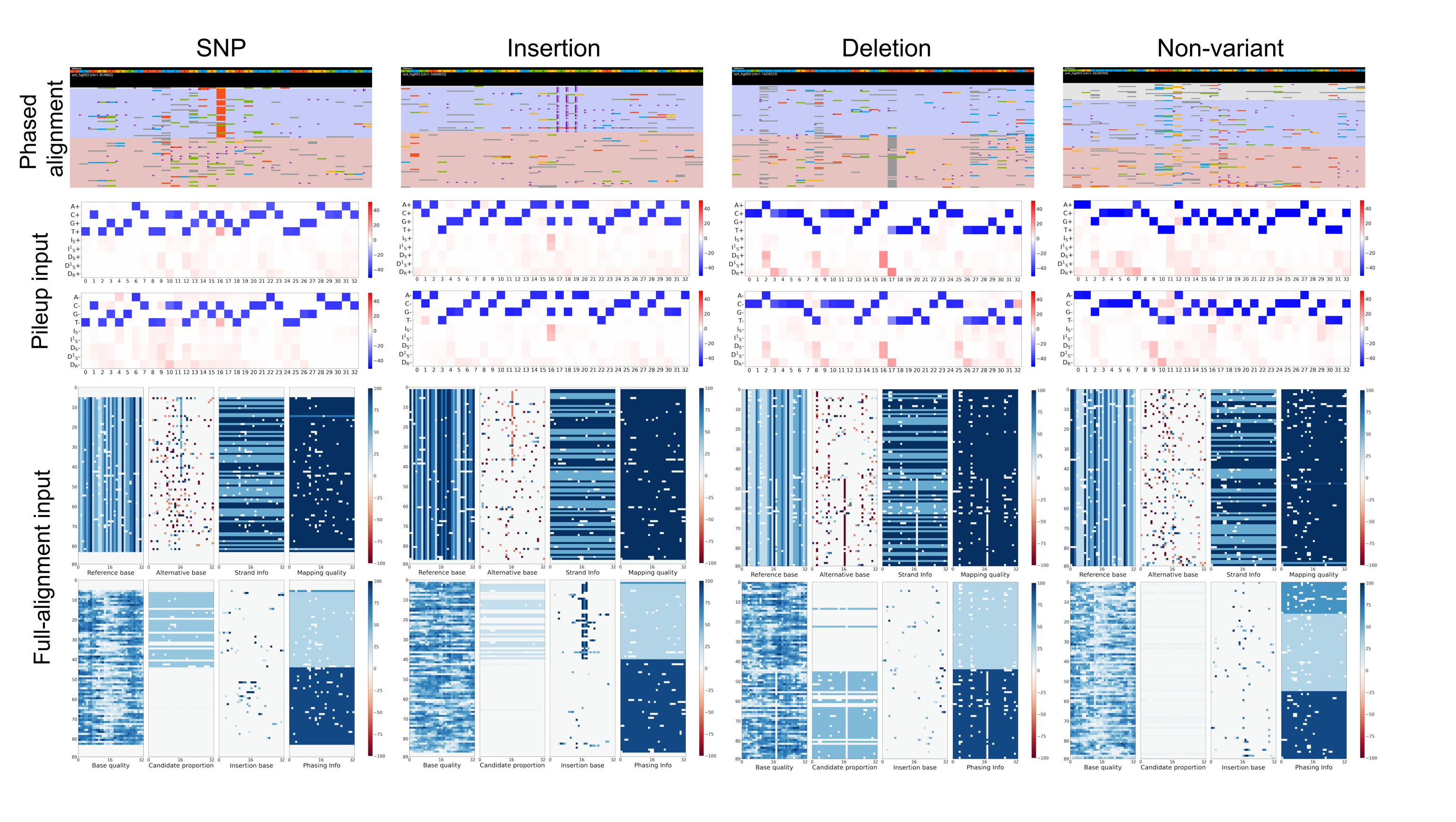

#### **Supplementary Figure 2.** Network input visualization.

The figure shows how the phased alignment, pileup input, and full-alignment input look like for a random SNP, insertion, deletion, and non-variant. In the phased alignment, A, C, G, T, insertion, and deletion are in green, blue, yellow, red, purple dots, and grey, respectively.

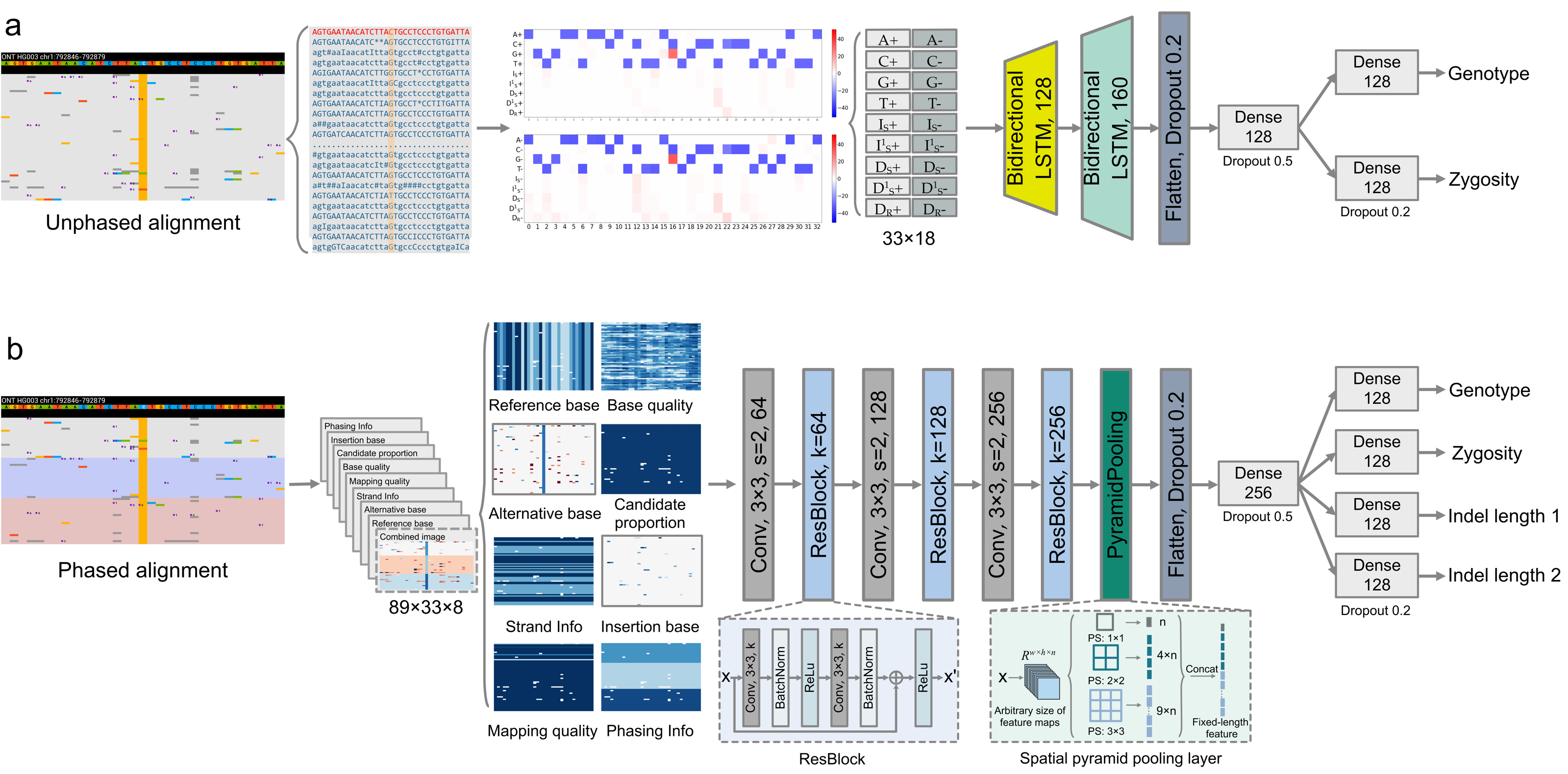

#### **Supplementary Figure 3.** Clair3 neural network architectures.

(a) The pileup network architecture. (b) The full-alignment network architecture.

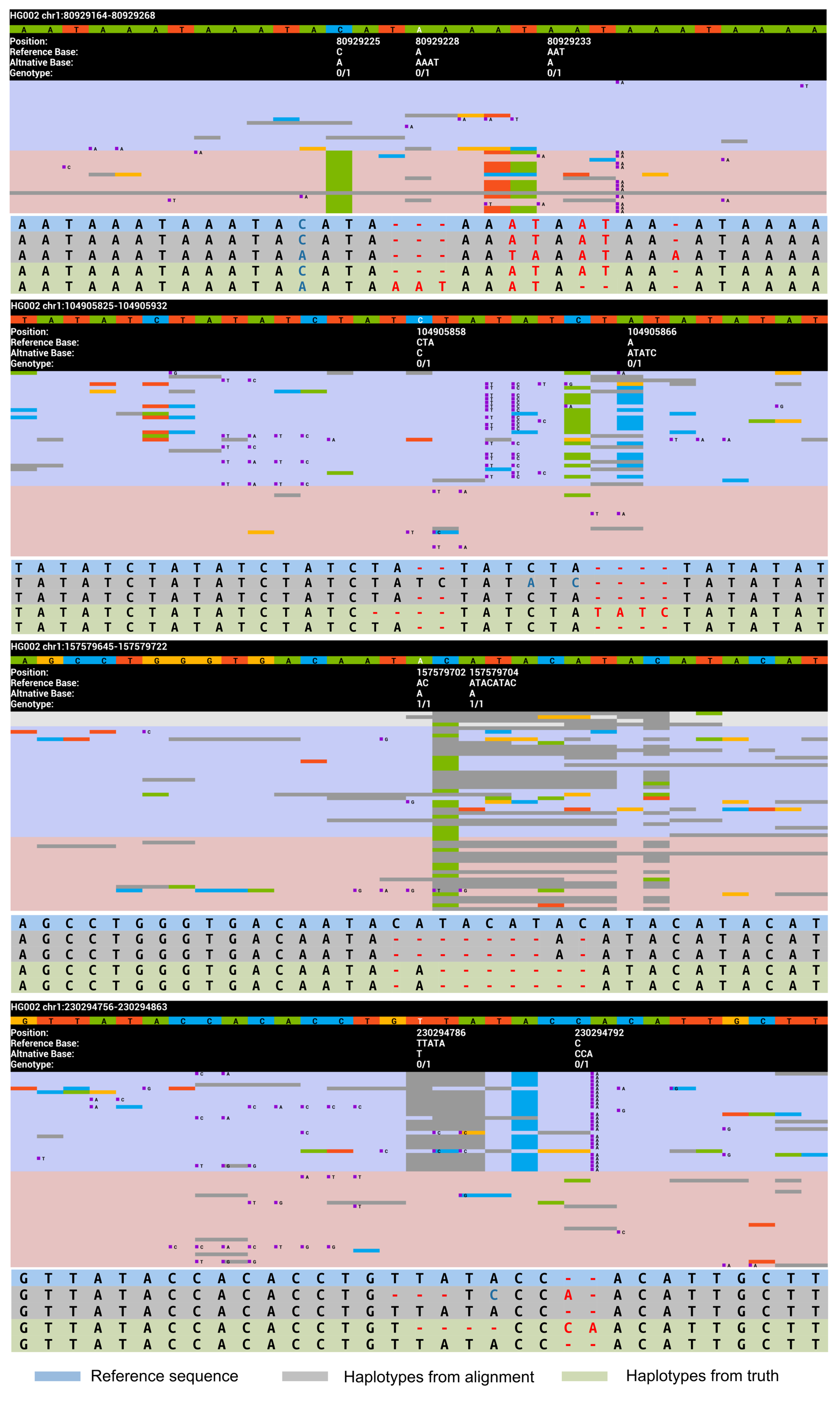

#### **Supplementary Figure 4.** Cases requiring representation unification.

The four cases show how the alignment can be discrepant with the truth, thus requiring representation unification (RU) before model training. The alignments (upper) and the haplotypes (lower) are shown for each case. In the alignments, the A, C, G, T, insertion, and deletion are in green, blue, yellow, red, purple dots, and grey, respectively. The truth details are shown in the second sub-caption. In the haplotypes, blue means SNP; red means Indel. DeepVariant has a module called “HaplotypeLabeler” that does a similar job, but only on cases with at least an exact match between the candidates and truth. Representation unification in Clair3, however, searches every truth and makes changes to the candidates even when no candidate has a direct match with the truth. In noisy long-reads, it happens more frequently to find no candidate directly matching the truth

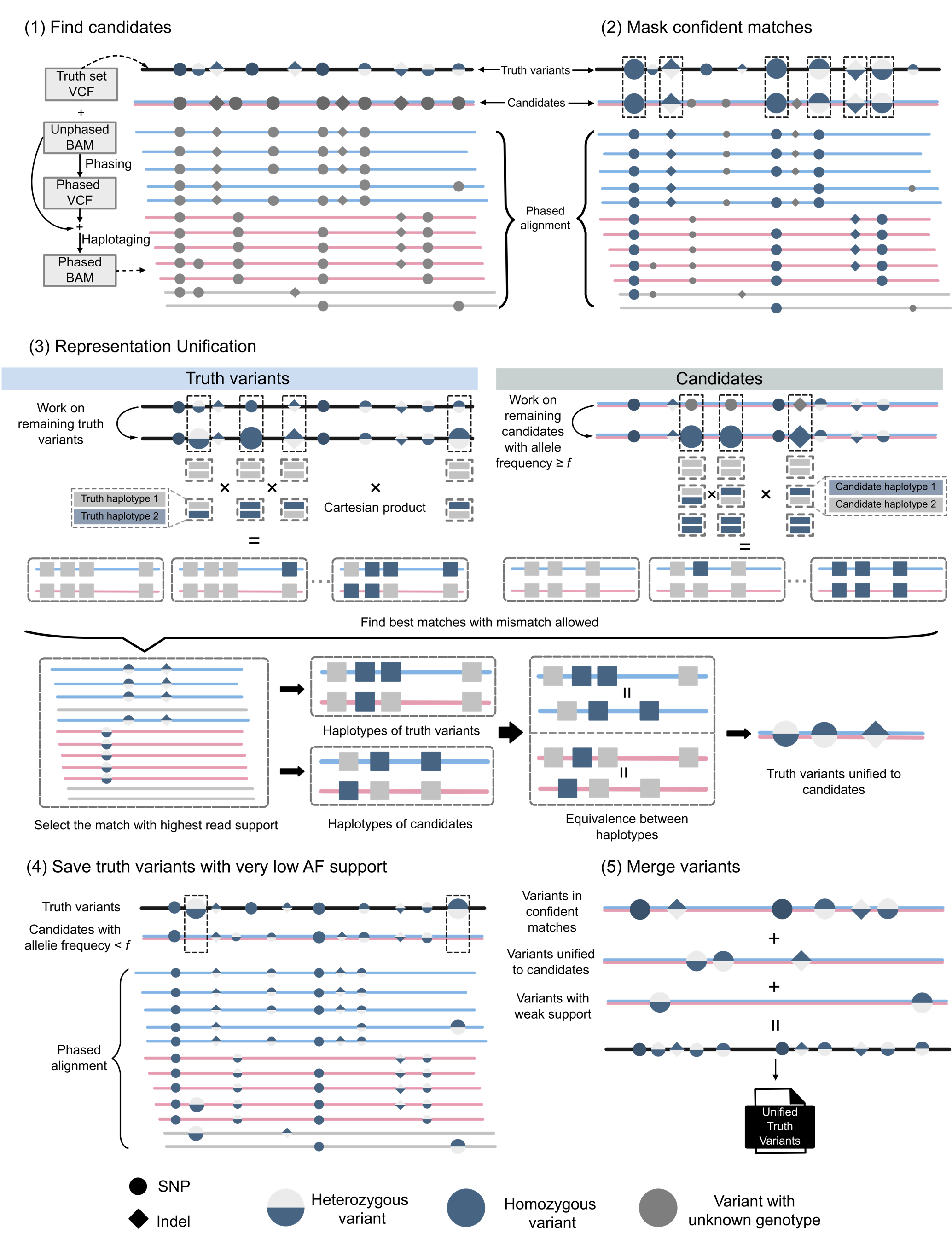

#### **Supplementary Figure 5.** The workflow for representation unification.

The detail of each step is described in the Methods section in the maintext.

### Supplementary Tables

#### **Supplementary Table 1.** A summary of the datasets used for training and testing.

Benchmarking settings (used in the last two columns): 1: HG003 for testing and other samples for training. 2: HG004 for testing and other samples for training. The model used to benchmark HG003, contained absolutely no known truths from HG003, and the model used to benchmark HG004, contained absolutely no known truths from HG004. For benchmarks on Guppy5 data, the performance of HG003 was tested on a model trained with HG002, HG004, and HG005, the performance of HG004 was tested on a model trained with HG002, HG003, and HG005. For benchmarks on Guppy4 data, the performance of HG003 was tested on a model trained with HG001, HG002, HG004, and HG005, the performance of HG004 was tested on a model trained with HG001, HG002, HG003, and HG005.

| **Platform and version** | **Sample** | **Reference** | **Aligner** | **Coverage** | **Source** | **Basecaller** | **Excluded chr20 from training** | **Used in training** | **Used in testing** |
| --- | --- | --- | --- | --- | --- | --- | --- | --- | --- |
| ONT Guppy 5 | HG002 | GRCh38_no_alt | minimap2 | 70.00 | EPI2ME labs | Guppy v5.0.6 | ✓ | 1,2 |  |
|  | HG002 | GRCh38_no_alt | minimap2 | 117.37 | HPRC | Guppy v5.0.14 | ✓ | 1,2 |  |
|  | HG003 | GRCh38_no_alt | minimap2 | 77.10 | HPRC | Guppy v5.0.14 | ✓ | 2 | 1 |
|  | HG004 | GRCh38_no_alt | minimap2 | 79.04 | HPRC | Guppy v5.0.14 | ✓ | 1 | 2 |
|  | HG005 | GRCh38_no_alt | minimap2 | 39.04 | HPRC | Guppy v5.0.14 | ✓ | 1,2 |  |
| ONT Guppy 4 | HG001 | GRCh38_no_alt | minimap2 | 58.1 | HPRC (Circulomics) | Guppy v4.2.2 | ✓ | 1,2 |  |
|  | HG001 | GRCh38_no_alt | minimap2 | 35.3 | HPRC (NBT) | Guppy v4.2.2 | ✓ | 1,2 |  |
|  | HG002 | GRCh38_no_alt | minimap2 | 432.4 | HPRC | Guppy v4.2.2 | ✓ | 1,2 |  |
|  | HG002 | GRCh38_no_alt | minimap2 | 49.0 | precisionFDA | Guppy v3.6.0 | ✓ | 1,2 |  |
|  | HG003 | GRCh38_no_alt | minimap2 | 85.0 | HPRC | Guppy v4.2.2 | ✓ | 2 | 1 |
|  | HG003 | GRCh38_no_alt | minimap2 | 84.8 | precisionFDA | Guppy v3.6.0 | ✓ | 2 |  |
|  | HG004 | GRCh38_no_alt | minimap2 | 87.5 | HPRC | Guppy v4.2.2 | ✓ | 1 | 2 |
|  | HG004 | GRCh38_no_alt | minimap2 | 87.4 | precisionFDA | Guppy v3.6.0 | ✓ | 1 |  |
|  | HG005 | GRCh38_no_alt | minimap2 | 57.0 | HPRC | Guppy v4.2.2 | ✓ | 1,2 |  |

#### **Supplementary Table 2.** HG003 multiple-coverage benchmarking results using Guppy 5 data.

| **Coverage** | **Caller** | **Overall** | | | **SNP** | | | **Indel** | | | **Insertion** | | | **Deletion** | | |
| --- | --- | --- | --- | --- | --- | --- | --- | --- | --- | --- | --- | --- | --- | --- | --- | --- |
|  |  | **Precision** | **Recall** | **F1-score** | **Precision** | **Recall** | **F1-score** | **Precision** | **Recall** | **F1-score** | **Precision** | **Recall** | **F1-score** | **Precision** | **Recall** | **F1-score** |
| 10x | Clair3 | 96.01% | 90.21% | 93.02% | 97.11% | 96.53% | 96.82% | 83.76% | 48.51% | 61.44% | 84.59% | 48.91% | 61.98% | 83.01% | 48.16% | 60.95% |
|  | PEPPER | 97.37% | 85.61% | 91.11% | 98.50% | 91.83% | 95.05% | 84.44% | 44.58% | 58.35% | 85.68% | 45.88% | 59.76% | 83.29% | 43.40% | 57.07% |
|  | Medaka | 47.36% | 87.74% | 61.52% | 52.99% | 94.86% | 67.99% | 18.24% | 40.83% | 25.22% | 11.29% | 42.94% | 17.88% | 47.07% | 38.91% | 42.61% |
|  | Clair | 84.01% | 66.81% | 74.43% | 84.38% | 73.84% | 78.76% | 76.11% | 20.48% | 32.28% | 69.73% | 20.68% | 31.90% | 83.24% | 20.31% | 32.65% |
| 20x | Clair3 | 98.17% | 94.54% | 96.32% | 99.22% | 99.42% | 99.32% | 88.56% | 62.33% | 73.17% | 88.49% | 62.74% | 73.42% | 88.63% | 61.97% | 72.94% |
|  | PEPPER | 98.26% | 94.28% | 96.23% | 99.66% | 98.99% | 99.32% | 86.06% | 63.26% | 72.92% | 87.88% | 62.80% | 73.25% | 84.51% | 63.68% | 72.63% |
|  | Medaka | 93.68% | 93.09% | 93.38% | 96.64% | 98.77% | 97.69% | 69.37% | 55.67% | 61.77% | 73.65% | 54.48% | 62.63% | 66.04% | 56.75% | 61.04% |
|  | Clair | 92.77% | 86.47% | 89.51% | 93.99% | 94.73% | 94.36% | 74.14% | 31.99% | 44.69% | 64.17% | 32.41% | 43.07% | 86.84% | 31.61% | 46.35% |
| 30x | Clair3 | 98.56% | 95.55% | 97.03% | 99.49% | 99.65% | 99.57% | 90.63% | 68.46% | 78.00% | 89.98% | 69.23% | 78.25% | 91.25% | 67.76% | 77.77% |
|  | PEPPER | 98.41% | 95.67% | 97.02% | 99.74% | 99.53% | 99.64% | 87.71% | 70.22% | 78.00% | 89.38% | 69.26% | 78.04% | 86.29% | 71.10% | 77.96% |
|  | Medaka | 96.56% | 94.08% | 95.30% | 98.89% | 99.12% | 99.01% | 77.40% | 60.84% | 68.13% | 82.68% | 59.14% | 68.96% | 73.39% | 62.38% | 67.43% |
|  | Clair | 94.58% | 89.14% | 91.78% | 96.28% | 97.26% | 96.77% | 71.89% | 35.58% | 47.60% | 60.48% | 35.92% | 45.07% | 87.29% | 35.27% | 50.24% |
| 40x | Clair3 | 98.73% | 96.11% | 97.40% | 99.60% | 99.70% | 99.65% | 91.71% | 72.39% | 80.91% | 90.70% | 73.58% | 81.25% | 92.67% | 71.31% | 80.60% |
|  | PEPPER | 98.50% | 96.35% | 97.41% | 99.77% | 99.65% | 99.71% | 88.87% | 74.58% | 81.10% | 90.37% | 73.50% | 81.07% | 87.60% | 75.55% | 81.13% |
|  | Medaka | 97.22% | 94.53% | 95.85% | 99.22% | 99.21% | 99.21% | 80.90% | 63.63% | 71.23% | 85.35% | 61.77% | 71.67% | 77.44% | 65.32% | 70.87% |
|  | Clair | 95.24% | 89.91% | 92.50% | 97.30% | 97.89% | 97.59% | 69.94% | 37.31% | 48.66% | 57.62% | 37.58% | 45.49% | 87.32% | 37.07% | 52.05% |
| 50x | Clair3 | 98.80% | 96.52% | 97.64% | 99.63% | 99.73% | 99.68% | 92.19% | 75.32% | 82.91% | 91.11% | 76.86% | 83.38% | 93.24% | 73.93% | 82.47% |
|  | PEPPER | 98.59% | 96.76% | 97.66% | 99.77% | 99.69% | 99.73% | 89.76% | 77.44% | 83.15% | 90.86% | 76.49% | 83.06% | 88.82% | 78.30% | 83.23% |
|  | Medaka | 97.51% | 94.80% | 96.13% | 99.29% | 99.25% | 99.27% | 83.00% | 65.40% | 73.16% | 86.77% | 63.50% | 73.33% | 80.04% | 67.13% | 73.02% |
|  | Clair | 95.50% | 90.23% | 92.79% | 97.85% | 98.10% | 97.98% | 68.15% | 38.32% | 49.06% | 55.15% | 38.55% | 45.38% | 87.25% | 38.11% | 53.05% |

#### **Supplementary Table 3.** HG004 multiple-coverage benchmarking results using Guppy 5 data.

| **Coverage** | **Caller** | **Overall** | | | **SNP** | | | **Indel** | | | **Insertion** | | | **Deletion** | | |
| --- | --- | --- | --- | --- | --- | --- | --- | --- | --- | --- | --- | --- | --- | --- | --- | --- |
|  |  | **Precision** | **Recall** | **F1-score** | **Precision** | **Recall** | **F1-score** | **Precision** | **Recall** | **F1-score** | **Precision** | **Recall** | **F1-score** | **Precision** | **Recall** | **F1-score** |
| 10x | Clair3 | 95.81% | 89.71% | 92.66% | 96.97% | 96.23% | 96.60% | 82.80% | 46.94% | 59.92% | 84.15% | 47.11% | 60.40% | 81.59% | 46.80% | 59.48% |
|  | PEPPER | 97.23% | 84.58% | 90.47% | 98.37% | 90.96% | 94.52% | 83.92% | 42.81% | 56.70% | 85.66% | 43.84% | 58.00% | 82.34% | 41.88% | 55.52% |
|  | Medaka | 43.82% | 87.06% | 58.30% | 49.31% | 94.37% | 64.77% | 16.08% | 39.15% | 22.80% | 9.90% | 41.00% | 15.95% | 41.77% | 37.47% | 39.51% |
|  | Clair | 82.32% | 65.42% | 72.90% | 82.74% | 72.41% | 77.23% | 73.19% | 19.61% | 30.93% | 66.91% | 19.74% | 30.48% | 80.17% | 19.49% | 31.36% |
| 20x | Clair3 | 98.19% | 94.40% | 96.25% | 99.33% | 99.49% | 99.41% | 87.66% | 60.99% | 71.93% | 88.25% | 60.86% | 72.04% | 87.12% | 61.11% | 71.83% |
|  | PEPPER | 98.22% | 94.07% | 96.10% | 99.70% | 98.99% | 99.34% | 85.25% | 61.82% | 71.67% | 87.82% | 60.77% | 71.83% | 83.13% | 62.77% | 71.53% |
|  | Medaka | 92.41% | 92.82% | 92.61% | 95.94% | 98.72% | 97.31% | 64.61% | 54.16% | 58.92% | 70.62% | 52.53% | 60.25% | 60.24% | 55.62% | 57.84% |
|  | Clair | 91.77% | 85.94% | 88.76% | 93.12% | 94.29% | 93.70% | 71.43% | 31.21% | 43.44% | 61.36% | 31.50% | 41.63% | 84.34% | 30.94% | 45.27% |
| 30x | Clair3 | 98.57% | 95.48% | 97.00% | 99.56% | 99.75% | 99.65% | 90.09% | 67.54% | 77.20% | 90.05% | 67.64% | 77.25% | 90.13% | 67.45% | 77.16% |
|  | PEPPER | 98.38% | 95.58% | 96.96% | 99.79% | 99.61% | 99.70% | 87.09% | 69.20% | 77.12% | 89.62% | 67.43% | 76.96% | 85.02% | 70.80% | 77.26% |
|  | Medaka | 95.94% | 93.87% | 94.89% | 98.77% | 99.12% | 98.94% | 73.48% | 59.47% | 65.74% | 81.23% | 57.11% | 67.07% | 68.05% | 61.61% | 64.67% |
|  | Clair | 93.83% | 88.94% | 91.32% | 95.70% | 97.17% | 96.43% | 69.32% | 35.01% | 46.52% | 57.74% | 35.26% | 43.78% | 85.15% | 34.78% | 49.39% |
| 40x | Clair3 | 98.77% | 96.10% | 97.42% | 99.65% | 99.81% | 99.73% | 91.52% | 71.81% | 80.47% | 91.20% | 72.21% | 80.60% | 91.81% | 71.44% | 80.36% |
|  | PEPPER | 98.53% | 96.33% | 97.42% | 99.85% | 99.76% | 99.80% | 88.48% | 73.85% | 80.51% | 90.87% | 71.96% | 80.32% | 86.53% | 75.57% | 80.68% |
|  | Medaka | 96.72% | 94.34% | 95.51% | 99.12% | 99.22% | 99.17% | 77.54% | 62.30% | 69.09% | 84.16% | 59.64% | 69.81% | 72.79% | 64.70% | 68.50% |
|  | Clair | 94.60% | 89.81% | 92.14% | 96.86% | 97.89% | 97.37% | 67.46% | 36.82% | 47.64% | 55.01% | 36.99% | 44.23% | 85.25% | 36.68% | 51.29% |
| 50x | Clair3 | 98.85% | 96.54% | 97.68% | 99.70% | 99.83% | 99.77% | 92.18% | 74.97% | 82.69% | 91.58% | 75.51% | 82.77% | 92.73% | 74.48% | 82.61% |
|  | PEPPER | 98.65% | 96.81% | 97.72% | 99.86% | 99.81% | 99.84% | 89.68% | 77.12% | 82.93% | 91.60% | 75.21% | 82.60% | 88.10% | 78.84% | 83.21% |
|  | Medaka | 97.14% | 94.60% | 95.85% | 99.28% | 99.26% | 99.27% | 80.05% | 64.01% | 71.14% | 85.83% | 61.24% | 71.48% | 75.82% | 66.51% | 70.86% |
|  | Clair | 94.94% | 90.16% | 92.49% | 97.51% | 98.14% | 97.82% | 65.76% | 37.87% | 48.06% | 52.71% | 38.00% | 44.17% | 85.18% | 37.75% | 52.31% |

#### **Supplementary Table 4.** HG003 multiple-coverage benchmarking results using Guppy 4 data.

| **Coverage** | **Caller** | **Overall** | | | **SNP** | | | **Indel** | | | **Insertion** | | | **Deletion** | | |
| --- | --- | --- | --- | --- | --- | --- | --- | --- | --- | --- | --- | --- | --- | --- | --- | --- |
|  |  | **Precision** | **Recall** | **F1-score** | **Precision** | **Recall** | **F1-score** | **Precision** | **Recall** | **F1-score** | **Precision** | **Recall** | **F1-score** | **Precision** | **Recall** | **F1-score** |
| 10x | Clair3 | 88.96% | 86.11% | 87.51% | 90.56% | 93.35% | 91.93% | 69.59% | 38.33% | 49.43% | 70.82% | 39.04% | 50.33% | 68.47% | 37.68% | 48.61% |
|  | PEPPER | 55.80% | 84.60% | 67.25% | 56.68% | 92.09% | 70.17% | 44.16% | 35.16% | 39.15% | 38.48% | 38.04% | 38.26% | 52.30% | 32.54% | 40.12% |
|  | Medaka | 27.29% | 84.41% | 41.25% | 30.94% | 92.48% | 46.36% | 8.35% | 31.18% | 13.17% | 5.04% | 35.39% | 8.83% | 35.69% | 27.35% | 30.97% |
|  | Clair | 80.32% | 59.67% | 68.47% | 80.54% | 66.22% | 72.68% | 74.85% | 16.47% | 27.00% | 75.18% | 16.17% | 26.62% | 74.57% | 16.74% | 27.34% |
| 20x | Clair3 | 97.47% | 92.79% | 95.07% | 99.17% | 98.77% | 98.97% | 80.91% | 53.33% | 64.29% | 80.11% | 54.10% | 64.59% | 81.68% | 52.64% | 64.02% |
|  | PEPPER | 88.98% | 92.31% | 90.62% | 90.40% | 98.95% | 94.48% | 73.73% | 48.53% | 58.53% | 70.17% | 51.66% | 59.51% | 77.75% | 45.70% | 57.57% |
|  | Medaka | 89.26% | 91.69% | 90.46% | 92.65% | 98.39% | 95.43% | 60.00% | 47.56% | 53.06% | 56.07% | 50.09% | 52.91% | 64.55% | 45.26% | 53.21% |
|  | Clair | 92.86% | 83.94% | 88.17% | 93.70% | 92.23% | 92.96% | 78.28% | 29.21% | 42.55% | 73.17% | 29.41% | 41.96% | 83.74% | 29.03% | 43.12% |
| 30x | Clair3 | 98.13% | 94.10% | 96.07% | 99.50% | 99.40% | 99.45% | 85.40% | 59.18% | 69.91% | 84.70% | 59.93% | 70.20% | 86.07% | 58.50% | 69.66% |
|  | PEPPER | 97.18% | 93.38% | 95.24% | 98.71% | 99.46% | 99.08% | 81.89% | 53.25% | 64.54% | 78.85% | 56.52% | 65.84% | 85.24% | 50.30% | 63.26% |
|  | Medaka | 95.43% | 93.09% | 94.25% | 98.40% | 98.99% | 98.69% | 70.45% | 54.15% | 61.23% | 68.77% | 56.42% | 61.98% | 72.20% | 52.09% | 60.51% |
|  | Clair | 95.62% | 88.18% | 91.75% | 96.70% | 96.39% | 96.55% | 79.24% | 33.99% | 47.57% | 72.28% | 34.29% | 46.51% | 87.10% | 33.71% | 48.61% |
| 40x | Clair3 | 98.62% | 94.63% | 96.58% | 99.61% | 99.55% | 99.58% | 89.47% | 62.16% | 73.36% | 88.86% | 62.86% | 73.63% | 90.06% | 61.52% | 73.10% |
|  | PEPPER | 98.20% | 93.82% | 95.96% | 99.48% | 99.58% | 99.53% | 85.49% | 55.84% | 67.55% | 82.84% | 58.94% | 68.88% | 88.33% | 53.02% | 66.27% |
|  | Medaka | 96.33% | 93.66% | 94.98% | 98.92% | 99.15% | 99.03% | 74.67% | 57.43% | 64.92% | 72.59% | 59.68% | 65.50% | 76.81% | 55.40% | 64.37% |
|  | Clair | 96.57% | 89.43% | 92.86% | 97.81% | 97.48% | 97.64% | 79.05% | 36.32% | 49.77% | 70.93% | 36.61% | 48.29% | 88.52% | 36.05% | 51.24% |
| 50x | Clair3 | 98.79% | 95.01% | 96.86% | 99.67% | 99.60% | 99.63% | 90.86% | 64.73% | 75.60% | 90.55% | 65.29% | 75.87% | 91.16% | 64.23% | 75.36% |
|  | PEPPER | 98.50% | 94.07% | 96.24% | 99.61% | 99.63% | 99.62% | 87.62% | 57.42% | 69.38% | 85.53% | 60.40% | 70.80% | 89.81% | 54.72% | 68.00% |
|  | Medaka | 96.71% | 93.97% | 95.32% | 99.00% | 99.22% | 99.11% | 77.42% | 59.33% | 67.18% | 75.25% | 61.69% | 67.80% | 79.66% | 57.19% | 66.58% |
|  | Clair | 97.09% | 89.91% | 93.37% | 98.23% | 97.92% | 98.07% | 81.00% | 37.09% | 50.88% | 72.72% | 36.96% | 49.01% | 90.41% | 37.20% | 52.71% |

#### **Supplementary Table 5.** HG004 multiple-coverage benchmarking results using Guppy 4 data.

| **Coverage** | **Caller** | **Overall** | | | **SNP** | | | **Indel** | | | **Insertion** | | | **Deletion** | | |
| --- | --- | --- | --- | --- | --- | --- | --- | --- | --- | --- | --- | --- | --- | --- | --- | --- |
|  |  | **Precision** | **Recall** | **F1-score** | **Precision** | **Recall** | **F1-score** | **Precision** | **Recall** | **F1-score** | **Precision** | **Recall** | **F1-score** | **Precision** | **Recall** | **F1-score** |
| 10x | Clair3 | 86.73% | 86.48% | 86.60% | 88.52% | 94.00% | 91.18% | 65.14% | 37.14% | 47.31% | 64.71% | 37.90% | 47.81% | 65.55% | 36.46% | 46.86% |
|  | PEPPER | 52.67% | 83.80% | 64.69% | 53.56% | 91.47% | 67.56% | 40.72% | 33.55% | 36.79% | 34.77% | 36.50% | 35.62% | 49.76% | 30.88% | 38.11% |
|  | Medaka | 25.67% | 83.71% | 39.29% | 29.35% | 91.93% | 44.50% | 7.33% | 29.78% | 11.76% | 4.40% | 33.96% | 7.79% | 33.53% | 26.00% | 29.29% |
|  | Clair | 79.11% | 58.24% | 67.09% | 79.36% | 64.72% | 71.30% | 72.85% | 15.70% | 25.83% | 72.61% | 15.40% | 25.41% | 73.06% | 15.97% | 26.20% |
| 20x | Clair3 | 97.74% | 92.57% | 95.08% | 98.89% | 99.08% | 98.98% | 85.15% | 49.93% | 62.94% | 84.27% | 50.82% | 63.40% | 85.98% | 49.12% | 62.52% |
|  | PEPPER | 87.25% | 92.07% | 89.60% | 88.74% | 98.92% | 93.56% | 71.10% | 47.12% | 56.68% | 66.60% | 50.31% | 57.32% | 76.40% | 44.23% | 56.03% |
|  | Medaka | 88.04% | 91.40% | 89.69% | 91.85% | 98.34% | 94.99% | 56.00% | 45.88% | 50.44% | 51.19% | 48.46% | 49.79% | 61.84% | 43.54% | 51.10% |
|  | Clair | 92.48% | 83.26% | 87.63% | 93.24% | 91.70% | 92.46% | 78.86% | 27.90% | 41.22% | 74.11% | 27.70% | 40.33% | 83.70% | 28.09% | 42.06% |
| 30x | Clair3 | 98.38% | 93.82% | 96.04% | 99.53% | 99.53% | 99.53% | 86.99% | 56.38% | 68.42% | 85.64% | 57.32% | 68.68% | 88.29% | 55.52% | 68.18% |
|  | PEPPER | 96.81% | 93.20% | 94.97% | 98.49% | 99.50% | 99.00% | 79.97% | 51.88% | 62.93% | 75.80% | 55.06% | 63.79% | 84.69% | 49.00% | 62.08% |
|  | Medaka | 94.81% | 92.84% | 93.82% | 98.31% | 99.01% | 98.66% | 66.27% | 52.40% | 58.53% | 62.98% | 54.60% | 58.49% | 69.83% | 50.42% | 58.56% |
|  | Clair | 95.46% | 87.74% | 91.44% | 96.41% | 96.18% | 96.29% | 80.21% | 32.42% | 46.17% | 73.63% | 32.20% | 44.80% | 87.26% | 32.62% | 47.49% |
| 40x | Clair3 | 98.59% | 94.43% | 96.47% | 99.65% | 99.63% | 99.64% | 88.63% | 60.30% | 71.77% | 86.84% | 61.47% | 71.99% | 90.38% | 59.24% | 71.57% |
|  | PEPPER | 98.06% | 93.66% | 95.81% | 99.49% | 99.63% | 99.56% | 83.89% | 54.49% | 66.07% | 80.33% | 57.67% | 67.14% | 87.81% | 51.61% | 65.01% |
|  | Medaka | 95.78% | 93.45% | 94.60% | 98.89% | 99.18% | 99.04% | 70.57% | 55.82% | 62.33% | 66.80% | 58.15% | 62.18% | 74.70% | 53.71% | 62.49% |
|  | Clair | 96.46% | 89.15% | 92.66% | 97.55% | 97.43% | 97.49% | 80.11% | 34.83% | 48.55% | 72.51% | 34.63% | 46.87% | 88.53% | 35.01% | 50.18% |
| 50x | Clair3 | 98.81% | 94.76% | 96.75% | 99.74% | 99.67% | 99.70% | 90.31% | 62.61% | 73.95% | 88.85% | 63.66% | 74.18% | 91.71% | 61.66% | 73.74% |
|  | PEPPER | 98.42% | 93.94% | 96.13% | 99.66% | 99.70% | 99.68% | 86.22% | 56.17% | 68.03% | 82.97% | 59.17% | 69.08% | 89.74% | 53.46% | 67.01% |
|  | Medaka | 96.29% | 93.76% | 95.01% | 99.10% | 99.27% | 99.18% | 73.43% | 57.70% | 64.62% | 69.19% | 60.12% | 64.33% | 78.13% | 55.51% | 64.90% |
|  | Clair | 96.93% | 89.69% | 93.17% | 98.13% | 97.86% | 98.00% | 79.77% | 36.17% | 49.77% | 71.44% | 35.94% | 47.82% | 89.19% | 36.38% | 51.68% |

#### **Supplementary Table 6.** Clair3’s performance by genomic context on HG003 at 50x of Guppy 4 data.

The stratification types and subtypes are from GIAB Genome Stratifications v2.0.

| **Stratification type** | **Stratification subtype** | **SNP** | | | | | | **Indel** | | | | | | |
| --- | --- | --- | --- | --- | --- | --- | --- | --- | --- | --- | --- | --- | --- | --- |
|  |  | **Precision** | **Recall** | **F1-Score** | **FP** | **FN** | **TP** | | **Precision** | **Recall** | **F1-Score** | **FP** | **FN** | **TP** |
| LowComplexity | Homopol_4-6bp | 99.60% | 99.48% | 99.54% | 3,304 | 4,353 | 824,853 | | 92.27% | 73.23% | 81.65% | 8,461 | 36,410 | 99,581 |
| LowComplexity | Homopol_7-11bp | 96.90% | 95.45% | 96.17% | 2,160 | 3,218 | 67,519 | | 79.89% | 46.82% | 59.04% | 12,498 | 56,234 | 49,518 |
| LowComplexity | Homopol_gt11bp | 89.79% | 83.97% | 86.78% | 1,169 | 1,953 | 10,234 | | 57.29% | 10.10% | 17.18% | 8,476 | 101,056 | 11,359 |
| LowComplexity | Imp_Homopol_gt10bp | 95.11% | 92.47% | 93.77% | 1,850 | 2,923 | 35,889 | | 68.92% | 18.70% | 29.42% | 13,032 | 124,485 | 28,639 |
| LowComplexity | TR_lt51bp | 98.69% | 98.47% | 98.58% | 540 | 627 | 40,379 | | 91.02% | 86.90% | 88.91% | 8,192 | 11,330 | 75,178 |
| LowComplexity | TR51-220bp | 99.08% | 99.08% | 99.08% | 246 | 244 | 26,257 | | 91.02% | 82.98% | 86.81% | 2,507 | 4,708 | 22,948 |
| LowComplexity | TR201-10kbp | 99.83% | 99.76% | 99.80% | 17 | 24 | 10,018 | | 96.86% | 90.97% | 93.82% | 120 | 357 | 3,595 |
| LowComplexity | TR_gt100bp | 99.46% | 99.49% | 99.47% | 108 | 103 | 19,927 | | 93.93% | 85.42% | 89.47% | 709 | 1,786 | 10,461 |
| LowComplexity | TR_and_Homopol | 97.63% | 96.55% | 97.09% | 3,919 | 5,734 | 160,618 | | 84.04% | 46.83% | 60.15% | 30,173 | 170,172 | 149,900 |
| SegmentalDuplications | ChainSelf | 97.86% | 98.67% | 98.26% | 3,501 | 2,152 | 159,830 | | 91.14% | 68.76% | 78.38% | 1,362 | 6,248 | 13,753 |
| SegmentalDuplications | ChainSelf_gt10kb | 94.94% | 97.21% | 96.06% | 3,101 | 1,667 | 58,172 | | 91.28% | 83.34% | 87.13% | 413 | 854 | 4,272 |
| SegmentalDuplications | SegDups | 96.57% | 98.02% | 97.29% | 4,245 | 2,411 | 119,549 | | 93.33% | 86.52% | 89.79% | 637 | 1,371 | 8,796 |
| SegmentalDuplications | SegDups_gt10kb | 96.07% | 97.74% | 96.90% | 4,217 | 2,385 | 102,976 | | 93.02% | 86.31% | 89.54% | 570 | 1,188 | 7,488 |
| Mappability | LowMap | 98.10% | 98.70% | 98.40% | 3,688 | 2,496 | 190,025 | | 94.29% | 88.15% | 91.12% | 535 | 1,185 | 8,815 |
| OtherDiffcult | L1H | 99.94% | 99.74% | 99.84% | 3 | 14 | 5,389 | | 98.51% | 93.43% | 95.90% | 3 | 14 | 199 |
| OtherDiffcult | MHC | 99.69% | 99.37% | 99.53% | 61 | 124 | 19,419 | | 91.84% | 82.41% | 86.87% | 131 | 301 | 1,410 |
| FunctionalRegions | CDS | 99.70% | 99.54% | 99.62% | 62 | 96 | 20,586 | | 87.09% | 84.05% | 85.54% | 47 | 59 | 311 |

#### **Supplementary Table 7.** HG001-HG006 multiple-coverage benchmarking results of chromosome 20 only.

HG001 has only Guppy4 fastq available provided by HPRC. No fast5 of HG001 is available thus we could not do basecalling using the Guppy5 basecaller on fast5 like what we have done on HG002-7.

| **Testing Dataset** | **Coverage** | **Caller** | **Overall** | | | **SNP** | | | **Indel** | | |
| --- | --- | --- | --- | --- | --- | --- | --- | --- | --- | --- | --- |
|  |  |  | **Precision** | **Recall** | **F1-score** | **Precision** | **Recall** | **F1-score** | **Precision** | **Recall** | **F1-score** |
| HG001 chr20 Guppy4 data | 10 | Clair3 | 95.30% | 82.94% | 88.69% | 96.45% | 90.15% | 93.20% | 77.77% | 32.96% | 46.29% |
|  | 10 | PEPPER | 49.25% | 81.52% | 61.41% | 50.06% | 88.99% | 64.07% | 36.94% | 29.73% | 32.94% |
|  | 20 | Clair3 | 98.07% | 91.52% | 94.69% | 99.42% | 97.92% | 98.67% | 82.18% | 47.15% | 59.92% |
|  | 20 | PEPPER | 84.85% | 91.55% | 88.07% | 86.43% | 98.41% | 92.03% | 66.21% | 43.97% | 52.85% |
|  | 30 | Clair3 | 98.63% | 93.48% | 95.98% | 99.71% | 99.35% | 99.53% | 86.52% | 52.80% | 65.58% |
|  | 30 | PEPPER | 96.30% | 93.02% | 94.63% | 98.22% | 99.38% | 98.80% | 75.67% | 48.91% | 59.41% |
| HG002 chr20 Guppy5 data | 10 | Clair3 | 95.50% | 88.93% | 92.10% | 96.74% | 95.90% | 96.32% | 81.41% | 44.74% | 57.75% |
|  | 10 | PEPPER | 96.99% | 83.03% | 89.47% | 98.21% | 89.64% | 93.73% | 83.04% | 41.14% | 55.02% |
|  | 20 | Clair3 | 98.24% | 93.80% | 95.97% | 99.50% | 99.46% | 99.48% | 86.49% | 57.96% | 69.41% |
|  | 20 | PEPPER | 98.16% | 93.46% | 95.75% | 99.76% | 98.86% | 99.31% | 84.21% | 59.25% | 69.56% |
|  | 30 | Clair3 | 98.61% | 95.04% | 96.79% | 99.71% | 99.82% | 99.77% | 89.19% | 64.74% | 75.02% |
|  | 30 | PEPPER | 98.27% | 95.09% | 96.65% | 99.82% | 99.64% | 99.73% | 85.83% | 66.25% | 74.78% |
| HG003 chr20 Guppy5 data | 10 | Clair3 | 96.53% | 90.42% | 93.38% | 97.74% | 96.92% | 97.33% | 82.97% | 47.47% | 60.39% |
|  | 10 | PEPPER | 97.49% | 86.23% | 91.52% | 98.72% | 92.59% | 95.56% | 83.37% | 44.24% | 57.81% |
|  | 20 | Clair3 | 98.38% | 94.39% | 96.35% | 99.50% | 99.55% | 99.52% | 87.91% | 60.37% | 71.58% |
|  | 20 | PEPPER | 98.26% | 94.25% | 96.21% | 99.77% | 99.22% | 99.49% | 84.92% | 61.46% | 71.31% |
|  | 30 | Clair3 | 98.68% | 95.33% | 96.97% | 99.64% | 99.73% | 99.68% | 90.24% | 66.27% | 76.42% |
|  | 30 | PEPPER | 98.39% | 95.54% | 96.94% | 99.79% | 99.67% | 99.73% | 86.95% | 68.29% | 76.50% |
| HG004 chr20 Guppy5 data | 10 | Clair3 | 96.39% | 89.66% | 92.90% | 97.63% | 96.43% | 97.03% | 82.21% | 45.55% | 58.63% |
|  | 10 | PEPPER | 97.54% | 84.85% | 90.76% | 98.77% | 91.46% | 94.97% | 83.15% | 41.85% | 55.67% |
|  | 20 | Clair3 | 98.40% | 94.06% | 96.18% | 99.59% | 99.48% | 99.53% | 87.12% | 58.78% | 70.20% |
|  | 20 | PEPPER | 98.29% | 93.76% | 95.97% | 99.80% | 99.03% | 99.41% | 84.80% | 59.43% | 69.88% |
|  | 30 | Clair3 | 98.70% | 95.13% | 96.88% | 99.76% | 99.76% | 99.76% | 89.44% | 64.95% | 75.25% |
|  | 30 | PEPPER | 98.36% | 95.18% | 96.74% | 99.88% | 99.68% | 99.78% | 85.82% | 65.87% | 74.53% |
| HG005 chr20 Guppy5 data | 10 | Clair3 | 95.71% | 91.98% | 93.81% | 96.65% | 96.47% | 96.56% | 84.23% | 55.57% | 66.96% |
|  | 10 | PEPPER | 97.32% | 86.63% | 91.66% | 98.17% | 91.02% | 94.46% | 86.67% | 51.00% | 64.22% |
|  | 20 | Clair3 | 98.58% | 96.40% | 97.48% | 99.55% | 99.53% | 99.54% | 88.94% | 70.94% | 78.93% |
|  | 20 | PEPPER | 98.84% | 95.90% | 97.35% | 99.79% | 98.99% | 99.39% | 89.43% | 70.80% | 79.03% |
|  | 30 | Clair3 | 98.93% | 97.36% | 98.14% | 99.74% | 99.87% | 99.80% | 91.29% | 76.99% | 83.53% |
|  | 30 | PEPPER | 98.97% | 97.30% | 98.13% | 99.80% | 99.74% | 99.77% | 91.24% | 77.57% | 83.85% |
| HG006 chr20 Guppy5 data | 10 | Clair3 | 95.87% | 91.58% | 93.67% | 96.83% | 96.62% | 96.73% | 84.12% | 52.67% | 64.78% |
|  | 10 | PEPPER | 97.34% | 86.33% | 91.50% | 98.28% | 91.38% | 94.71% | 85.27% | 47.38% | 60.92% |
|  | 20 | Clair3 | 98.56% | 95.78% | 97.15% | 99.56% | 99.65% | 99.61% | 88.38% | 65.87% | 75.48% |
|  | 20 | PEPPER | 98.62% | 95.43% | 97.00% | 99.75% | 99.22% | 99.48% | 87.43% | 66.18% | 75.34% |
|  | 30 | Clair3 | 98.84% | 96.63% | 97.72% | 99.71% | 99.85% | 99.78% | 90.55% | 71.78% | 80.08% |
|  | 30 | PEPPER | 98.70% | 96.62% | 97.65% | 99.83% | 99.75% | 99.79% | 88.30% | 72.49% | 79.62% |

#### **Supplementary Table 8.** Benchmarking results on different pileup and full-alignment workload distributions.

| **Sample** | **Runtime (mins)** | **Proportion of low-quality pileup calls that enters full-alignment calling (variant / reference)** | **Overall** | | | **SNP** | | | **Indel** | | | **Insertion** | | | **Deletion** | | |
| --- | --- | --- | --- | --- | --- | --- | --- | --- | --- | --- | --- | --- | --- | --- | --- | --- | --- |
|  |  |  | **Precision** | **Recall** | **F1** | **Precision** | **Recall** | **F1** | **Precision** | **Recall** | **F1** | **Precision** | **Recall** | **F1** | **Precision** | **Recall** | **F1** |
| ONT HG003 50x of Guppy 5 data | 81 | 0.0/0.0 (pileup only) | 95.60% | 95.36% | 95.48% | 99.40% | 99.65% | 99.53% | 70.01% | 67.06% | 68.50% | 70.55% | 68.67% | 69.60% | 69.50% | 65.61% | 67.50% |
|  | 185 | 0.1/0.1 | 97.82% | 96.23% | 97.02% | 99.47% | 99.75% | 99.61% | 85.33% | 73.03% | 78.70% | 84.26% | 74.33% | 78.99% | 86.37% | 71.85% | 78.44% |
|  | 190 | 0.2/0.1 | 98.54% | 96.40% | 97.46% | 99.59% | 99.74% | 99.66% | 90.35% | 74.39% | 81.60% | 89.11% | 75.76% | 81.89% | 91.55% | 73.16% | 81.33% |
|  | 194 | 0.3/0.1 (default) | 98.75% | 96.50% | 97.61% | 99.63% | 99.73% | 99.68% | 91.88% | 75.16% | 82.68% | 90.75% | 76.68% | 83.13% | 92.97% | 73.77% | 82.27% |
|  | 200 | 0.4/0.1 | 98.78% | 96.51% | 97.64% | 99.63% | 99.73% | 99.68% | 92.11% | 75.28% | 82.85% | 91.01% | 76.83% | 83.32% | 93.17% | 73.87% | 82.41% |
|  | 205 | 0.5/0.1 | 98.79% | 96.51% | 97.64% | 99.63% | 99.73% | 99.68% | 92.16% | 75.30% | 82.88% | 91.07% | 76.85% | 83.36% | 93.21% | 73.90% | 82.44% |
|  | 210 | 0.6/0.1 | 98.79% | 96.52% | 97.64% | 99.63% | 99.73% | 99.68% | 92.18% | 75.31% | 82.89% | 91.10% | 76.86% | 83.37% | 93.22% | 73.91% | 82.45% |
|  | 216 | 0.7/0.1 | 98.80% | 96.52% | 97.64% | 99.63% | 99.73% | 99.68% | 92.20% | 75.32% | 82.91% | 91.12% | 76.86% | 83.38% | 93.25% | 73.93% | 82.47% |
|  | 222 | 0.8/0.1 | 98.80% | 96.52% | 97.64% | 99.63% | 99.73% | 99.68% | 92.22% | 75.33% | 82.92% | 91.14% | 76.86% | 83.40% | 93.27% | 73.94% | 82.49% |
|  | 227 | 0.9/0.1 | 98.80% | 96.52% | 97.65% | 99.63% | 99.73% | 99.68% | 92.23% | 75.33% | 82.93% | 91.15% | 76.86% | 83.40% | 93.27% | 73.95% | 82.50% |
|  | 232 | 1.0/0.1 | 98.80% | 96.52% | 97.65% | 99.63% | 99.73% | 99.68% | 92.24% | 75.34% | 82.94% | 91.16% | 76.86% | 83.40% | 93.28% | 73.95% | 82.50% |
|  | 313 | 1.0/0.3 | 98.79% | 96.53% | 97.65% | 99.63% | 99.73% | 99.68% | 92.21% | 75.43% | 82.98% | 91.13% | 76.92% | 83.42% | 93.26% | 74.09% | 82.58% |
|  | 582 | 1.0/1.0 (full-alignment only) | 98.79% | 96.53% | 97.65% | 99.63% | 99.73% | 99.68% | 92.21% | 75.44% | 82.99% | 91.13% | 76.92% | 83.42% | 93.26% | 74.10% | 82.58% |

#### **Supplementary Table 9.** Network performance with channels or tasks removed.

(a) results of the insertion, phasing, MQ, or BQ channel being removed (b) results of the two Indel length tasks being removed.

**(a)**

| **Testing  dataset** | **Setting** | **Overall** | | | **SNP** | | | **Indel** | | |
| --- | --- | --- | --- | --- | --- | --- | --- | --- | --- | --- |
|  |  | **Precision** | **Recall** | **F1-score** | **Precision** | **Recall** | **F1-score** | **Precision** | **Recall** | **F1-score** |
| HG003  20X of Guppy5 data | Clair3 | 98.17% | 94.54% | 96.32% | 99.22% | 99.42% | 99.32% | 88.56% | 62.33% | 73.17% |
|  | w/o insertion channel | 98.15% | 94.53% | 96.31% | 99.21% | 99.42% | 99.31% | 88.42% | 62.31% | 73.11% |
|  | w/o phasing channel | 98.13% | 93.28% | 95.64% | 99.06% | 99.19% | 99.13% | 88.33% | 54.28% | 67.24% |
|  | w/o MQ channel | 97.92% | 92.52% | 95.14% | 99.06% | 99.39% | 99.22% | 84.63% | 47.22% | 60.62% |
|  | w/o BQ channel | 88.38% | 91.17% | 89.76% | 93.57% | 98.91% | 96.17% | 46.86% | 40.10% | 43.22% |

**(b)**

| **Testing  dataset** | **Setting** | **Overall** | | | **SNP** | | | **Indel** | | |
| --- | --- | --- | --- | --- | --- | --- | --- | --- | --- | --- |
|  |  | **Precision** | **Recall** | **F1-score** | **Precision** | **Recall** | **F1-score** | **Precision** | **Recall** | **F1-score** |
| HG003  20X of Guppy5 data | Default | 98.17% | 94.54% | 96.32% | 99.22% | 99.42% | 99.32% | 88.56% | 62.33% | 73.17% |
|  | Removing two Indel length tasks | 94.44% | 94.94% | 94.69% | 99.27% | 99.32% | 99.30% | 64.08% | 66.05% | 65.05% |

#### **Supplementary Table 10.** Memory consumption of Clair3, PEPPER, Clair, and Medaka when using 24 cores.

| **Testing  dataset** | **Caller** | **Peak memory** | **Average  memory** | **Peak memory  per core** | **Average memory  per core** |
| --- | --- | --- | --- | --- | --- |
| ONT HG003 20X of Guppy5 data | Clair3 | 11GB | 7.8GB | 0.46GB | 0.33GB |
|  | PEPPER | 89GB | 62GB | 3.71GB | 2.58GB |
|  | Clair | 18GB | 13GB | 0.75GB | 0.54GB |
|  | Medaka | 6GB | 2.8GB | 0.25GB | 0.12GB |

### Supplementary Notes

#### Description of full-alignment input channels

- **Reference base:** An integer {A:100, G:75, T:50, C:25} is assigned to a position according to the reference base. 0 is assigned to deleted bases.
- **Alternative base:** An integer {A:100, G:75, T:50, C:25, insertion starting position: -50, deletion starting position: -100, deletion non-starting positions: 0} is assigned to a position if the alignment at the position mismatches the reference base. 0 is assigned if a position matches the reference base.
- **Strand information:** An integer {+: 100, -:50} is assigned all positions of a read according to the strand the read aligned to. 0 is assigned if a position is in a deletion.
- **Mapping quality:** An integer ranging from 0 to 100 scaled up from the Phred mapping score (0 to 60, cap at 60 if >60) of an aligned read is assigned to the whole read. 0 is assigned to deleted bases. Decimal places are truncated.
- **Base quality:** An integer ranging from 0 to 100 scaled up from the Phred base quality score (0 to 40, cap at 40 if >40) is assigned to each aligned base. 0 is assigned to deleted bases. Decimal places are truncated.
- **Candidate proportion:** An integer ranging from 0 to 100 to indicate the percentage of reads among all reads that are supporting an exact mismatch pattern in a read is assigned to all positions of the read. 0 is assigned to deleted bases. For example, a candidate with 20 reads, 10 supporting the reference allele A, 5 supporting alternative allele C, 3 supporting alternative allele T, and 2 supporting an insertion AT, the integer assigned to the reads that support A, C, T, and AT are 0, 25, 15, and 10, respectively. Decimal places are truncated.
- **Insertion base:** Inserted bases are exhibited after each insertion’s starting position with integers {A:100, G:75, T:50, C:25}. Although rarely happens, it is possible for exhibited insertions to have overlaps. But an overlap can be disambiguated using the “alternative base” channel, in which the insertion starting positions are assigned with -50.
- **Phasing information:** For unphased reads, an integer 60 is assigned to all reads. For phased reads, {HP1: 30, unphased: 60, HP2: 90} is assigned to all positions of each read. 0 is assigned to deleted bases. If the reads are phased, the reads in all eight channels will be sorted in the order “unphased, HP1, HP2”.

#### Evaluation metrics

We used Illumina’s Haplotype Comparison Tools (aka. hap.py) available at <https://github.com/Illumina/hap.py> to benchmark our results and generated three metrics Precision, Recall, and F1-score for each of the three categories including Overall, SNP, and Indel, respectively. From the number of true positives (*TP*), false positives (*FP*), true negatives (*TN*), and false negatives (*FN*), *hap.py* computes the three metrics as Precision = $TP\div(TP+FP)$, Recall = $TP\div\left( TP+FN \right)$, and F1-score = $2TP/(2TP+FN+FP)$.

#### Network outputs of the pileup and full-alignment network

The output of both the pileup and full-alignment network has four tasks, including: 1) the 21-genotype probabilistic model (21 probabilities); 2) zygosity (3 probabilities); 3) the length of the first indel allele 33 probabilities); and 4) the length of the second indel allele (33 probabilities). There are 90 probabilities in total. The details of the four tasks were given in Clair’s manuscript, and are given here again for clarity.

The 21-genotype probabilistic model comprises all of the possible genotypes of a diploid sample at a genome position, including ‘AA’, ‘AC’, ‘AG’, ‘AT’, ‘CC’, ‘CG’, ‘CT’, ‘GG’, ‘GT’, ‘TT’, ‘AI’, ‘CI’, ‘GI’, ‘TI’, ‘AD’, ‘CD’, ‘GD’, ‘TD’, ‘II’, ‘DD’, and ‘ID’, where ‘A’, ‘C’, ‘G’, ‘T’, ‘I’ (insertion) and ‘D’ (deletion) denote the six possible alleles. The zygosity task outputs the probability of the input being 1) a homozygous reference (0/0); 2) heterozygous with 1 or 2 alternative alleles (0/1 or 1/2); or 3) a homozygous variant (1/1). The zygosity task is partially redundant to the 21-genotype task, but it makes decisions independently, and it crosschecks the decision made by the 21-genotype task. Tasks three and four have the same design. They output the length of up to two indel alleles. Each task outputs 33 probabilities, including the likelihood of 1) more than 15bp deleted (<−15bp); 2) any number between −15bp and 15bp, including 0bp; and 3) more than 15bp inserted (>15bp). In training, the indel allele with a smaller number is set as the first indel allele. For example, for a heterozygous 1bp deletion, the first indel allele is set as −1bp and the second as 0bp (−1bp/0bp). For a heterozygous 1-bp insertion, 0bp/1bp is set. This design makes the non-0-bp training variants for both tasks balanced. For a heterozygous indel with two alternative alleles, say, one −2bp and one 5bp, −2bp/5bp is set. For a homozygous indel, two indel alleles are set to the same value. For indels longer than 15bp, the exact length is determined directly from the CIGAR strings of the supporting reads. The output of the two indel allele tasks are also used for crosschecking with the 21-genotype task, with 0bp supporting an SNP allele, and non-0bp supporting an indel allele.

#### Training data augmentation using subsampled coverage

Lower coverage usually leads to lower precision and recall in variant calling. To train Clair3 to achieve better performance on variants with lower coverages, we subsampled each dataset into four or nine additional datasets with lower coverages. We subsampled each dataset into nine additional datasets with lower coverages. The subsampling factors *f* are calculated as $f={(\sqrt[h]{8/c})}^{n}$, where *c* is the average coverage of a sample, 8 is the minimum coverage after subsampling, *h* is either 4 or 9, and *n* is from 1 to *h*. Using HG002 as an example, its average coverage is 70.00x and the nine subsampled coverages are 55.09x, 43.23x, 33.97x, 26.69x, 20.97x, 16.48x, 12.95x, 10.18x and 8.00x. We used the command ‘samtools view -s *f*’ to generate subsampled BAM files.

#### Data sources

##### Reference genomes

###### GRCh38_no_alt

https://[ftp.ncbi.nlm.nih.gov/genomes/all/GCA/000/001/405/GCA_000001405.15_GRCh38/seqs_for_alignment_pipelines.ucsc_ids/GCA_000001405.15_GRCh38_no_alt_analysis_set.fna.gz](http://ftp.ncbi.nlm.nih.gov/genomes/all/GCA/000/001/405/GCA_000001405.15_GRCh38/seqs_for_alignment_pipelines.ucsc_ids/GCA_000001405.15_GRCh38_no_alt_analysis_set.fna.gz)

###### GRCh38 Stratification regions v2.0

<https://ftp-trace.ncbi.nlm.nih.gov/giab/ftp/release/genome-stratifications/v2.0/GRCh38/>

##### GIAB Truth Variants

###### HG001 (NA12878), GRCh38, v3.3.2

<https://ftp>[-trace.ncbi.nlm.nih.gov/giab/ftp/release/NA12878_HG001/NISTv3.3.2/GRCh38/](ftp://ftp-trace.ncbi.nlm.nih.gov/giab/ftp/release/NA12878_HG001/NISTv3.3.2/GRCh38/HG001_GRCh38_GIAB_highconf_CG-IllFB-IllGATKHC-Ion-10X-SOLID_CHROM1-X_v.3.3.2_highconf_PGandRTGphasetransfer.vcf.gz)

###### HG002 (NA24385), GRCh38, v4.2.1

https[://ftp-trace.ncbi.nlm.nih.gov/giab/ftp/release/AshkenazimTrio/HG002_NA24385_son/NISTv4.2.1/GRCh38/](ftp://ftp-trace.ncbi.nlm.nih.gov/giab/ftp/release/AshkenazimTrio/HG002_NA24385_son/NISTv4.2.1/GRCh38/)

###### HG003 (NA24149), GRCh38, v4.2.1

https[://ftp-trace.ncbi.nlm.nih.gov/giab/ftp/release/AshkenazimTrio/HG003_NA24149_father/NISTv4.2.1/GRCh38/](ftp://ftp-trace.ncbi.nlm.nih.gov/giab/ftp/release/AshkenazimTrio/HG003_NA24149_father/NISTv4.2.1/GRCh38/)

###### HG004 (NA24143), GRCh38, v4.2.1

https[://ftp-trace.ncbi.nlm.nih.gov/giab/ftp/release/AshkenazimTrio/HG004_NA24143_mother/NISTv4.2.1/GRCh38/](ftp://ftp-trace.ncbi.nlm.nih.gov/giab/ftp/release/AshkenazimTrio/HG004_NA24143_mother/NISTv4.2.1/GRCh38/)

###### HG005 (NA24631), GRCh38, v3.3.2

https[://ftp-trace.ncbi.nlm.nih.gov/giab/ftp/release/ChineseTrio/HG005_NA24631_son/NISTv3.3.2/GRCh38/](http://ftp-trace.ncbi.nlm.nih.gov/giab/ftp/release/ChineseTrio/HG005_NA24631_son/NISTv3.3.2/GRCh38/)

##### CMRG Truth Variants

###### HG002 (NA24385), GRCh38, v1.0.0

https://ftp-trace.ncbi.nlm.nih.gov/ReferenceSamples/giab/release/AshkenazimTrio/HG002_NA24385_son/CMRG_v1.00/GRCh38/SmallVariant/

##### Oxford Nanopore (ONT) Sequencing Data

###### HG002 Guppy 5.0.6 (NA24385), GRCh38_no_alt, 70.00-fold

https://labs.epi2me.io/gm24385_2021.05/

###### HG002 Guppy 5.0.14 (NA24385), GRCh38_no_alt, 117.37-fold

http://www.bio8.cs.hku.hk/guppy5_data/guppy5_sup/hg002/HG002_GM24385_guppy_v5.0.14_r9.4.1_sup_prom_117x.bam

###### HG003 Guppy 5.0.14 (NA24149), GRCh38_no_alt, 77.10-fold

http://www.bio8.cs.hku.hk/guppy5_data/guppy5_sup/hg003/HG003_GM24149_guppy_v5.0.14_r9.4.1_sup_prom_77x.bam

###### HG004 Guppy 5.0.14 (NA24143), GRCh38_no_alt, 79.04-fold

http://www.bio8.cs.hku.hk/guppy5_data/guppy5_sup/hg004/HG004_GM24143_guppy_v5.0.14_r9.4.1_sup_prom_79x.bam

###### HG005 Guppy 5.0.14 (NA24631), GRCh38_no_alt, 39.46-fold

http://www.bio8.cs.hku.hk/guppy5_data/guppy5_sup/hg005/HG005_GM24631_guppy_v5.0.14_r9.4.1_sup_prom_40x.bam

###### HG001 Circulomics Guppy 4.2.2 (NA12878), GRCh38_no_alt, 58.06-fold

<https://s3-us-west-2.amazonaws.com/human-pangenomics/index.html?prefix=NHGRI_UCSC_panel/HG001/nanopore/Guppy_4.2.2/HG001_Circulomics_Guppy_4.2.2.fastq.gz>

###### HG001 NBT Guppy 4.2.2 (NA12878), GRCh38_no_alt, 35.30-fold

<https://s3-us-west-2.amazonaws.com/human-pangenomics/index.html?prefix=NHGRI_UCSC_panel/HG001/nanopore/Guppy_4.2.2/HG001_NBT2018_Guppy_4.2.2.fastq.gz>

###### HG002 HD Guppy 4.2.2 (NA24385), GRCh38_no_alt, 432.38-fold

https://s3-us-west-2.amazonaws.com/human-pangenomics/index.html?prefix=NHGRI_UCSC_panel/HG002/nanopore/Guppy_4.2.2/

###### HG002 PrecisionFDA (NA24385), GRCh38_no_alt, 49.04-fold

<https://precision.fda.gov/files/file-Fpk3KGj0fQX2VjXjF8YXqz93-1>

###### HG003 Guppy 4.2.2 (NA24149), GRCh38_no_alt, 84.97-fold

https://s3-us-west-2.amazonaws.com/human-pangenomics/index.html?prefix=NHGRI_UCSC_panel/HG003/nanopore/Guppy_4.2.2/

###### HG003 PrecisionFDA (NA24149), GRCh38_no_alt, 84.80-fold

<https://precision.fda.gov/files/file-Fpk3VVQ0fQX12Pvb925XP843-1>

###### HG004 Guppy4.2.2 (NA24143), GRCh38_no_alt, 87.51-fold

[https://s3-us-west-2.amazonaws.com/human-pangenomics/index.html?prefix=NHGRI_UCSC_panel/HG004/nanopore/Guppy_4.2.2/](https://s3-us-west-2.amazonaws.com/human-pangenomics/index.html?prefix=NHGRI_UCSC_panel/HG004/nanopore/Guppy_4.2.2/GM24143_1-3_Guppy_4.2.2_prom.fastq.gz)

###### HG004 PrecisionFDA (NA24143), GRCh38_no_alt, 87.36-fold

<https://precision.fda.gov/files/file-Fpk3VZQ0fQXKP8xk381VGYvg-1>

###### HG005 Guppy 4.2.2 (NA24631), GRCh38_no_alt, 56.96-fold

[https://s3-us-west-2.amazonaws.com/human-pangenomics/index.html?prefix=NHGRI_UCSC_panel/HG005/nanopore/Guppy_4.2.2/](https://s3-us-west-2.amazonaws.com/human-pangenomics/index.html?prefix=NHGRI_UCSC_panel/HG005/nanopore/Guppy_4.2.2/01_09_20_R941_GM24631_1-3_Guppy_4.2.2_prom.fastq.gz)

#### Commands

##### Read alignment using Minimap2 (v2.17-r941)

```
# Align ONT reads to GRCh38_no_alt
minimap2 -t ${THREADS} -aL -z 600,200 -x map-ont ref.fa input.fastq.gz | samtools view -bh -o output.unsorted.bam -
samtools sort -@ ${THREADS} -o output.sorted.bam output.unsorted.bam
samtools index -@ ${THREADS} output.sorted.bam
```

##### BAM subsampling using Samtools (v1.10)

```
# Using ${FRAC} for both random seed and subsampling fraction
samtools view -@ ${THREADS} -s ${FRAC}.${FRAC} -b -o subsampled.bam ${BAM}
samtools index -@ ${THREADS} subsampled.bam
```

##### Coverage calculation using Mosdepth (v0.2.9)

mosdepth -t ${THREADS} -n -x --quantize 0:15:150: output ${BAM}

##### Clair3 model training

```
# For better versioning and easier update of the guides, we put the model training steps, commands, parameters, and caveats in the following three GitHub READMEs:

# Representation Unification:
https://github.com/HKU-BAL/Clair3/blob/main/docs/representation_unification.md

# Pileup model training:
https://github.com/HKU-BAL/Clair3/blob/main/docs/pileup_training.md

# Full-alignment model training:
https://github.com/HKU-BAL/Clair3/blob/main/docs/full_alignment_training.md
```

##### Running Clair3 (v0.1-r11)

```
docker run -it \
  -v ${INPUT_DIR}:${INPUT_DIR} \
  -v ${OUTPUT_DIR}:${OUTPUT_DIR} \
  hkubal/clair3:v0.1 \
  /opt/bin/run_clair3.sh \
  --bam_fn=${INPUT_DIR}/input.bam \
  --ref_fn=${INPUT_DIR}/ref.fa \
  --threads=${THREADS} \
  --platform="${PLATFORM}" \
  --model_path="/opt/models/${PLATFORM}" \
  --output=${OUTPUT_DIR}
```

##### Running PEPPER (r0.8)

```
docker run --ipc=host \
  -v "${INPUT_DIR}":"${INPUT_DIR}" \
  -v "${OUTPUT_DIR}":"${OUTPUT_DIR}" \
  kishwars/pepper_deepvariant:r0.8 \
  run_pepper_margin_deepvariant call_variant \
  -b "${BAM}" \
  -f "${REF}" \
  -o "${OUTPUT_DIR}" \
  -p "${SAMPLE}" \
  -t ${THREADS} \
  --ont_r9_guppy5_sup # --ont if using Guppy 4 data

Note: PEPPER has decided to drop support of Guppy 4 data since r0.5 https://github.com/kishwarshafin/pepper/issues/118. Therefore, for benchmarks using Guppy 4 data, PEPPER r0.4 was used.
```

##### Running Clair (v2.1.1)

```
# The ONT model was chosen from https://github.com/HKU-BAL/Clair
# Use quality cut-off 748 as suggested in https://github.com/HKU-BAL/Clair
python clair.py callVarBamParallel \
  --chkpnt_fn ${Clair_MODEL} \
  --ref_fn ${REF} \
  --bam_fn ${BAM} \
  --threshold 0.2 \
  --sampleName ${SAMPLE} \
  --pysam_for_all_indel_bases \
  --output_prefix "${OUTPUT_DIR}/${SAMPLE} > command.sh
cat command.sh | parallel -j${THREADS}

# concat parallel vcf output and filter LowQual variants
vcf-concat *.vcf | vcf-sort > output.vcf
bcftools view -i'QUAL>=748' -O z -o output_filterQual.vcf.gz output.vcf
```

##### Running Medaka (v1.4.4)

CHR_PREFIX='chr'

CHR=(1 2 3 4 5 6 7 8 9 10 11 12 13 14 15 16 17 18 19 20 21 22 X Y)

parallel -j1 medaka_variant -f ${REF} -i ${BAM} -t ${THREADS} -o ${OUTPUT_DIR}/{1} -r ${CHR_PREFIX}{1} ::: ${CHR[@]}

vcf-concat ${OUTPUT_DIR}/*/round_1.vcf | vcf-sort > ${OUTPUT_DIR}/medaka_output.vcf

Note: Medaka’s variant calling submodule was deprecated in v1.5.0. ONT recommended using Clair3 either directly or through the Oxford Nanopore Technologies provided Nextflow implementation available through EPI2ME Labs (<https://github.com/nanoporetech/medaka/blob/83ff752eda1b3da9c1d873fb9a62e89c13d3e657/README.md>).

##### Running Longshot (v0.4.5)

longshot --bam ${BAM} --ref ${REF} --out ${OUTPUT_DIR}/longshot_output.vcf

##### Benchmarking using hap.py (v0.3.12)

```
hap.py ${GIAB_BASELINE_VCF} output.vcf.gz \
  -o ${OUTPUT_DIR}/happy \
  -r ${REF} \
  -f ${GIAB_CONFIDENT_BED} \
  --threads ${THREADS} \
  --pass-only \
  --engine=vcfeval

# Get the Precision, Recall, F1-Score of insertions and deletions, respectively
pypy3 ${CLAIR3} GetOverallMetrics --happy_vcf_fn ${OUTPUT_DIR}/happy.vcf.gz --output_fn happy.log
```

##### Benchmarking using qfy.py (v0.3.12)

```
# Benchmarking all genome stratifications regions
qfy.py ${OUTPUT_DIR}/happy.vcf.gz \
  -t ga4gh \
  --stratification v2.0-GRCh38-stratifications.tsv \
  -o ${OUTPUT_PREFIX} \
  -r ${REF} \
  --threads ${THREADS}
```
